# Paclitaxel-loaded Cationic Fluid Lipid Nanodiscs and Liposomes with Brush-Conformation PEG Chains Penetrate Breast Tumors and Trigger Caspase-3 Activation

**DOI:** 10.1101/2022.03.28.486128

**Authors:** Lorena Simón-Gracia, Pablo Scodeller, William S. Fisher, Valeria Sidorenko, Victoria M. Steffes, Kai K. Ewert, Cyrus R. Safinya, Tambet Teesalu

## Abstract

Novel approaches are required to address the urgent need to develop lipid-based carriers of paclitaxel (PTX) and other hydrophobic drugs for cancer chemotherapy. Carriers based on cationic liposomes (CLs) with fluid (i.e., chain-melted) membranes (e.g., EndoTAG-1^®^) have shown promise in preclinical and late-stage clinical studies. Recent work found that the addition of a cone-shaped poly(ethylene glycol)-lipid (PEG-lipid) to PTX-loaded CLs (CLs_PTX_) promotes a transition to sterically stabilized, higher-curvature (smaller) nanoparticles consisting of a mixture of PEGylated CLs_PTX_ and PTX-containing fluid lipid nanodiscs (nanodiscs_PTX_). These CLs_PTX_ and nanodiscs_PTX_ show significantly improved uptake and cytotoxicity in cultured human cancer cells at PEG coverage in the brush regime (10 mol% PEG-lipid).

Here, we studied the PTX loading, *in vivo* circulation half-life, and biodistribution of systemically administered CLs_PTX_ and nanodiscs_PTX_ and assessed their ability to induce apoptosis in triple-negative breast cancer-bearing immunocompetent mice. We focused on *fluid* rather than *solid* lipid nanodiscs because of the significantly higher solubility of PTX in fluid membranes. At 5 and 10 mol% of a PEG-lipid (PEG5K-lipid, molecular weight of PEG 5000 g/mol), the mixture of PEGylated CLs_PTX_ and nanodiscs_PTX_ was able to incorporate up to 2.5 mol% PTX without crystallization for at least 20 h. Remarkably, compared to preparations containing 2 and 5 mol% PEG5K-lipid (with the PEG chains in the mushroom regime), the particles at 10 mol% (with PEG chains in the brush regime) showed significantly higher blood half-life, tumor penetration and proapoptotic activity. Our study suggests that increasing the PEG coverage of CL-based drug nanoformulations can improve their pharmacokinetics and therapeutic efficacy.

## Introduction

Many chemotherapeutic agents are poorly soluble in aqueous solutions and must be formulated with vectors (carriers) to enable their use. A prime example of this is paclitaxel (PTX) – a potent cytotoxic drug that targets tubulin to block mitotic spindle assembly, chromosome segregation, and cell division.^1–4^ PTX is highly hydrophobic with very low water solubility, and ineffective in its crystalline form. Nonetheless, PTX is one of the most commonly used anti-cancer drugs, with multi-billion dollar sales each year.^5–11^ Currently, the most widely used formulations of PTX are Taxol^®^(where PTX is solubilized using nonionic Kolliphor EL surfactant) and Abraxane (where PTX is formulated in albumin nanoparticles).^5,12–14^ Serious side effects, both due to vector-related toxicity and the indiscriminate delivery of PTX throughout the body, are common and dose-limiting for currently used PTX formulations.^15–17^ Efforts are therefore underway to develop improved PTX formulations with reduced side effects: novel carrier materials to avoid the hypersensitivity reactions associated with the use of Kolliphor EL (formerly Cremophor EL, polyoxyethylated castor oil), carriers with higher efficacy to allow administration of lower total doses of PTX, and precision-guided platforms to increase specificity of delivery of PTX to malignant tissues.^18–27^

Liposomes are widely used biocompatible carriers of payloads ranging from hydrophilic drugs and biologicals (nucleic acids and proteins) to hydrophobic drugs.^5,28–36^ A prominent example of a liposomal carrier of a hydrophobic drug is Doxil^®^,^37^ the formulation of another commonly used chemotherapeutic agent (doxorubicin). However, the chemical properties of doxorubicin enable efficient loading of the aqueous interior of the liposomes with the drug by using a transmembrane pH gradient – a process not transferable to most other drugs including PTX.^38^

Cationic liposomes [CLs; consisting of mixture of cationic (or ionizable) and neutral lipids] are prevalent nonviral vectors for the delivery of therapeutic nucleic acids (NAs), as recently demonstrated for the mRNA vaccines against COVID-19.^39–42^ In addition, CLs are suitable carriers for hydrophobic drugs. EndoTAG^®^-1, a cationic lipid formulation of PTX currently undergoing phase III clinical testing, is based on fluid-phase DOTAP/DOPC CLs with 3 mol% PTX.^19,25,35,43^ (DOTAP, *N*-[2,3-dioleoyloxy-1-propyl]-trimethylammonium chloride, is a univalent cationic lipid; DOPC, 1,2-dioleoyl-*sn*-glycerophospho-choline, is a naturally occurring neutral phospholipid.) EndoTAG-1 is taken up in endothelial cells in solid tumors via electrostatic interactions with cell surface anionic sulfated proteoglycans.^36,43–47^ To achieve PTX loading into liposomal membranes beyond the 3 mol% viewed as the maximal loading in most preclinical efficacy studies and clinical trials to date,^19,25,35,43,45^ the lipid tails that interact with PTX can be modified.^48^ Additionally, *in vivo* selectivity of systemic lipid carriers can be improved by conjugating the poly(ethylene glycol)-lipids (PEG-lipids) with affinity targeting ligands such as homing peptides.^49,50^

Modification of the lipids and the lipid composition can drive structural transitions of fluid-phase CL vectors,^29,51–55^ including the spontaneous formation of PEGylated fluid lipid micelles (discs, cylinders and spheroids) coexisting with vesicles.^56,57^ We recently reported that increasing the density of PEG-lipids strongly reduces the average size of DOTAP/DOPC CLs.^57^ Above a critical concentration near the mushroom–brush transition, the PEG-lipid component provides steric stabilization that suppresses the aggregation of nonPEGylated CLs and triggers the formation of nanometer-scale disc-shaped bicelles (fluid lipid nanodiscs). Figure 1 illustrates the effect of PEGylation on the morphology of CL-based vectors of PTX as revealed by cryogenic TEM. In a formulation with composition identical to that of EndoTAG-1 (Figure 1A), large vesicles of a variety of sizes coexist with smaller vesicles and occasional discs. In contrast, in a formulation that differs from EndoTAG-1 by replacement of a part of the neutral DOPC with PEG2K-lipid (to a total of 10 mol%), larger vesicles have disappeared completely and only an abundance of very small vesicles together with a large number of nanodiscs_PTX_ are observed.

**Figure 1.**
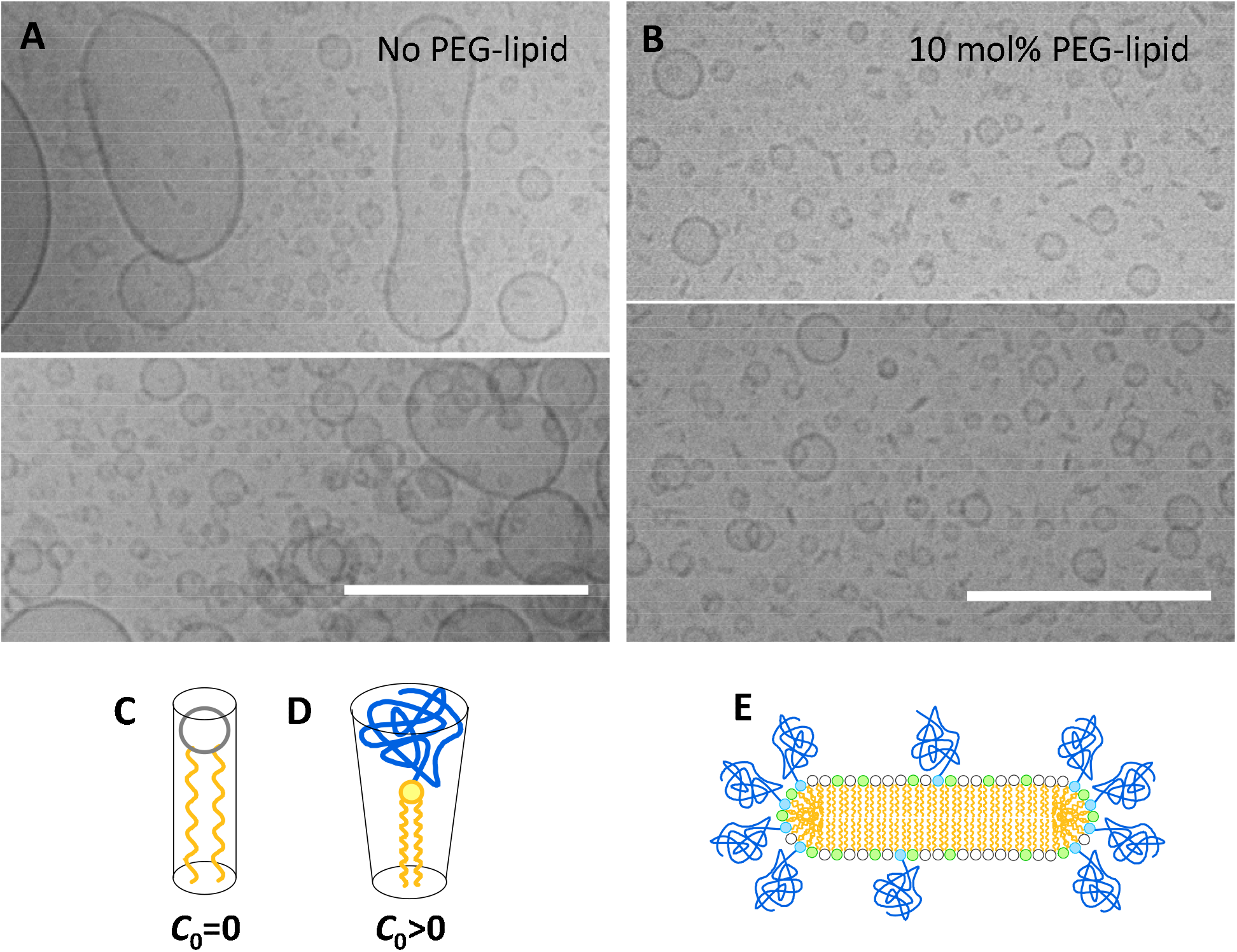
Cryogenic TEM images of PTX-loaded CLs with and without PEGylation, lipid shapes and curvature, and schematic of a lipid nanodisc. **(A)** Sonicated liposomes at the EndoTAG-1 composition (DOTAP/DOPC/PTX=50/47/3 molar ratio), exhibiting both larger and smaller vesicles with few discs. **(B)** Sonicated PTX-loaded CLs with 10 mol% PEG2K-lipid (DOTAP/DOPC/PEG2K-lipid/PTX=50/37/10/3 molar ratio), lacking vesicles above ~50 nm in size and showing a prevalence of very small vesicles and lipid discs. Scale bars: 200 nm. The images are previously unpublished micrographs of formulations investigated in ref. 57. See Figures S1–S4 in the Supporting Information for uncropped versions of the micrographs. **(C, D)** Two common shapes of lipid molecules which result in membrane spontaneous curvature *C*_0_=0 (**C**, e.g., DOPC; cylindrical shape yielding flat bilayers) and *C*_0_>0 (**D**, e.g., PEG-lipids; cone shape yielding micellar monolayer assemblies or high-curvature bilayers).^61–63^ **(E)** Schematic depiction of a disc micelle, with PEG-lipid segregated to the high-curvature edges.

Membrane physics of a multicomponent fluid lipid system provides a rationale for explaining a regime of coexistence of spherical vesicles and discoidal micelles. In our ternary lipid system, the majority components DOTAP and DOPC are cylindrically shaped lipids which prefer flat bilayer membranes with spontaneous curvature *C*_0_=0 (Figure 1C). The cone-shaped PEG-lipid, which prefers positive high curvature surfaces with *C*_0_>0 (Figure 1D), is the minority component. At low concentrations of PEG-lipid, with PEG chains in the mushroom conformation, the lipids remain mixed with maximal entropy. With increasing molar fraction of PEG-lipid near the brush regime (i.e. where PEG chains in the mushroom conformation begin to overlap), the formation of an alternative morphology, namely, lipid discs, begins to become increasingly favorable. Here, lipid phase separation and segregation of the cone-shaped PEG-lipids to the high curvature edges of discs takes place, reducing the curvature elastic energy cost of edge formation^58–60^ (i.e. compared to forming edges containing cylindrical shaped DOPC or DOTAP lipids) and stabilizing discs.

A number of previous studies have characterized the structure and properties of *solid* lipid nanodiscs.^64–67^ In these systems, the (charge-neutral or slightly anionic) lipid assemblies with saturated tails were studied in the chain-ordered “gel” phase with very low chain mobility, giving rise to 2D crystalline membranes.^68^ In our work, we focused on *fluid* lipid nanodiscs where the lipid tails contained one *cis* double bound, thus forming the chain-melted “fluid phase” with high chain mobility in the temperature range studied (between room temperature and ~37 °C). The main motivation for using fluid versus solid lipid nanodiscs is the significantly higher solubility of PTX in the nonpolar tail region of the membrane. Prior work has shown that membranes consisting of lipids containing one^69,72^ or two^48^ *cis* double bonds can accommodate much higher molar fractions of PTX than membranes consisting of lipids with saturated tails.^18,35,69–71^

Compared to formulations similar to the benchmark EndoTAG-1, the PEGylated lipid vectors consisting of mixtures of PEGylated CLsPTX and PTX-containing fluid lipid nanodiscs (nanodiscs_PTX_), with PEG chains in the brush state, show a significant PEG concentration-dependent enhancement of cellular uptake and cytotoxicity (i.e., efficacy).^57^ Thus, size, stabilization, and shape of the PEGylated lipid vectors have profound effects on their cellular uptake and payload delivery.

The observation that incorporating PEG-lipids yields fluid-phase CL carriers of PTX with stable micellar structures and improved *in vitro* efficacy prompted us to study the *in vivo* performance of systemically administered PEGylated PTX-loaded CLs. Here, we characterized the *in vivo* biodistribution and proapoptotic activity of a series of PEGylated CL vectors of PTX in triple-negative breast cancer (TNBC) mice. Compared to vectors with 2 and 5 mol% PEG5K-lipid, lipid nanoparticle vectors containing 10 mol% PEG5K-lipid possessed longer plasma circulation half-life and resulted in significantly improved TNBC accumulation, tumor extravasation, and activation of proapoptotic caspase-3.

## Materials and Methods

Samples for cryogenic TEM were prepared and imaged as described previously.^57^

### Materials

A stock solution of carboxyfluorescein-labeled DOPE (1,2-dioleoyl-*sn*-glycero-3-phosphoethanolamine-N-(carboxyfluorescein), DOPE-FAM) in chloroform was purchased from Avanti Polar Lipids. Other lipid stock solutions were prepared by dissolving DOPE-PEG5000, DOTAP, and DOPC, purchased from Avanti Polar Lipids, in chloroform at 3.64, 41.93, and 31.8 mM, respectively. A stock solution of PTX in chloroform was prepared by dissolving solid PTX (Biotang Inc.) in chloroform at 10 mM.

### Liposome preparation

Mixed solutions of lipid and PTX were prepared in chloroform:methanol (3:1, v/v) in small glass vials at 1 mM for PTX solubility, biodistribution, and caspase activation experiments and at 16.75 mM for lipid solubility experiments. The chloroform:methanol solvent was evaporated under a nitrogen stream for 10 min and the lipid was further dried in a vacuum for 16 h. The resultant film was resuspended in PBS to 1 mM for PTX solubility, biodistribution, and caspase activity experiments or 67 mM for lipid solubility experiments. This suspension was sonicated with a tip sonicator (30 W output) for 7 min.

### Differential Interference Contrast (DIC) microscopy

For PTX solubility experiments, 2 μL each of cationic lipid formulations containing 10 mol% PEG5K-lipid and 2, 2.5, or 3 mol% PTX were placed on glass microscope slides and covered by glass coverslips secured by parafilm cutouts. All remaining samples were incubated at room temperature (RT) for 20 h and 2 μL were again used for imaging. For lipid solubility experiments, 2 μL each of samples of formulations with 2, 5, and 10 mol% PEG5K-lipid were placed on glass microscope slides and again covered by coverslips secured by parafilm cutouts. All slides were imaged at 40× magnification on an inverted Ti2-E (Nikon) microscope.

### Determination of the half-life of PEGylated, PTX-containing CL vectors

The animal experiments were performed according to protocols approved by the Estonian Ministry of Agriculture, Committee of Animal Experimentation (projects #159 and #160). Eight week-old immunocompetent female BALB/c mice were intravenously (i.v.) injected with 100 μL of PEGylated cationic liposome formulations of PTX containing 2, 5, or 10 mol% PEG5K-lipid. After 15, 30, 60, 180, 360, and 1440 min circulation, 5 μL of blood was extracted from the tail vein and mixed with 50 μL of phosphate buffered saline (PBS, pH 7.4) with heparin at 4 °C. The samples were centrifuged at 300 *g* at 4 °C for 5 min. The fluorescence of the supernatant was measured at 490 nm/535 nm (excitation/emission) using a Victor X5 Multilabel Microplate Reader (Perkin Elmer, USA). The data were fitted to a curve using a biexponential decay formula to obtain the half-life of the PEGylated formulations using the Origin 2022b software.

### Detection of tumor accumulation of PEGylated PTX-containing CL vectors and cleaved caspase-3 immunostaining

TNBC cells (10^6^ 4T1 cells in 50 μL of PBS) were orthotopically injected into the mammary gland of 8 week-old immunocompetent female BALB/c mice. One week later, 100 μL of PEGylated CL formulations of PTX were i.v. administered into the tail vein, and 24 h later, the mice were anesthetized, perfused with PBS, and the tumor and organs were excised and kept in 4% paraformaldehyde (PFA) solution in PBS at 4 °C overnight. PFA-fixed tissues were washed and immersed in PBS at RT for 1 h.

Then, the tissues were incubated in 15% sucrose solution in PBS at 4 °C overnight. The following day, the 15% sucrose solution was replaced with 30% sucrose solution in PBS and incubated at 4 °C overnight. The cryoprotected tissues were frozen in OCT (optimal cutting temperature) compound and cryosectioned at 20 μm. The sections were air-dried at RT for 1 h, permeabilized with PBS containing 0.2% Triton-X for 15 min, washed with PBS containing 0.05% Tween-20 (PBST), and blocked with PBST containing 5% bovine serum albumin (BSA), 5% fetal bovine serum (FBS), and 5% goat serum for 1 h. The tissue sections were incubated at 4 °C overnight with rat anti-mouse CD31 (cat. no. 553370, BD Biosciences) and rabbit anti-cleaved Caspase-3 (cat. no. 9661, Cell Signaling Technology) as primary antibodies in blocking buffer diluted 1 in 5 in PBST (antibody dilution 1/200 and 1/400, respectively). Alexa 647-conjugated goat anti-rabbit IgG (cat. no. A21245, Thermo Fischer Scientific) and Alexa 546-conjugated goat anti-rat IgG (cat. no. A11081, Thermo Fischer Scientific) were used as secondary antibodies (antibody dilution 1/300). The slides were incubated with the secondary antibodies at RT for 2 h. The sections were washed with PBST and PBS, and the nuclei were stained with 1 μg/mL DAPI in PBS for 5 min. Stained slides were mounted with mounting medium and the coverslips were sealed with nail polish. The tissues were imaged using a fluorescence confocal microscope (FV1200MPE, Olympus), and the images analyzed using the Olympus FluoView Ver.4.2a Viewer program. To quantify the intensity of the FAM signal from PEGylated lipid vectors and cleaved caspase-3, confocal images were analyzed using ImageJ. Three to 6 random areas per tumor were chosen from 3 different planes and this was repeated for 3 tumors for each treatment group.

### Statistical tests

The statistical analyses were performed using the one-way ANOVA and Fisher LSD tests.

## Results and Discussion

### PTX membrane solubility in PEGylated fluid-phase CLs

Because PTX is incorporated into the hydrophobic environment formed by the lipid tails rather than in the aqueous interior of liposomes, the initial encapsulation efficiency in CLs is ≈100%. However, it is essential for the efficacy of PTX-loaded CLs that PTX remains soluble in the fluid membrane. We previously found that the membrane solubility of PTX is slightly lower in PEGylated CLs than in bare CLs.^57,72^ To determine the optimal PTX content for the CLs in the current study, we assessed the solubility of PTX in CLs containing 10 mol% PEG5K-lipid, the highest PEG-lipid content used. The formation of therapeutically inert PTX crystals, due to PTX self-association in the membrane and subsequent phase separation, is an indicator of PTX insolubility.^72,73^ Thus, we assessed PTX solubility in the PEGylated CL formulations by monitoring PTX crystal formation with DIC microscopy.^57,72^ Representative DIC micrographs of sonicated CLs containing 10 mol% PEG5K-lipid and 2 to 3 mol% PTX are displayed in Figure 2. Insoluble PTX formed crystals in the samples with 3 mol% PTX at RT after 20 h (Figure 2, top right panel, yellow arrow), whereas PTX remained soluble at 2 and 2.5 mol% (Figure 2, middle and bottom right panels). PTX remained soluble for days at 2 mol% PTX, which prompted us to choose this PTX loading for the biodistribution experiments.

**Figure 2.**
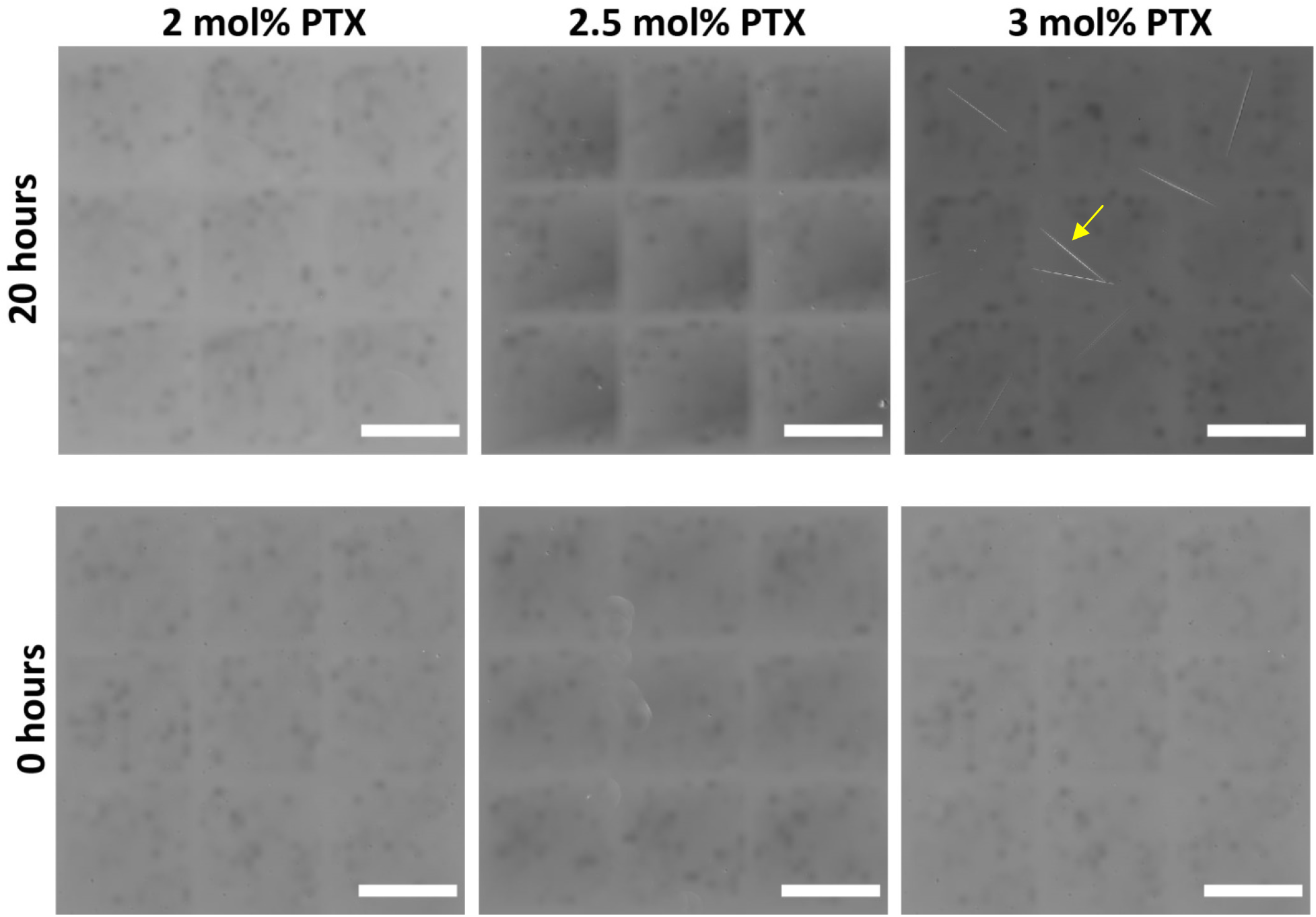
Solubility of PTX in PEGylated CLs at different drug loadings. DIC micrographs of sonicated CLs containing 10 mol% PEG5K-lipid, loaded with 2 (left column), 2.5 (center column), and 3 (right column) mol% PTX, imaged immediately (bottom row) and 20 h (top row) after resuspension in PBS. Whereas PTX crystals were not present in any of the samples immediately following resuspension (or at 20 h time point in the case of 2 or 2.5 mol% PTX), PTX crystal formation was evident 20 h after resuspension for CLs containing 3 mol% PTX (top right panel; the yellow arrow points to a representative crystal). CL composition: PEG5K-lipid:DOTAP:DOPC:PTX=10:50:40–x_PTX_:x_PTX_ molar ratio, with x_PTX_ the PTX content in mol%. Scale bars: 200 μm.

### PEGylation increases solubility of fluid-phase CLs at high lipid concentrations

The size distribution of nanoparticles has an important effect on pharmacokinetics and biodistribution, with a preferred particle size between 5 and 100 nm.^74^ At the low total lipid concentration used in the PTX solubility (Figure 2) and biodistribution (see below) studies, no particles were visible by DIC microscopy for any of the formulations investigated, meaning that the size of the lipid assemblies is below the diffraction limit of a few hundred nm. The small vesicles and discs visible in cryogenic TEM were undetectable by optical microscopy due to this reason. For systemic therapy, however, PTX-loaded CLs must be prepared at total lipid concentrations of ≥50 mg/mL to achieve sufficiently high PTX dosage.^75,76^ Therefore, we prepared sonicated CL suspensions with varying degrees of PEGylation at a total lipid concentration of 50 mg/mL. At this concentration, formulations with 2 mol% PEG5K-lipid contained spontaneously formed giant multilamellar structures, greater than ~100 μm in size, that were visible in DIC microscopy (Figure 3, left panel). These giant multilamellar structures are likely to be poorly suited for *in vivo* applications. However, such large structures were not found in CLs containing 5 and 10 mol% PEG5K-lipid (Figure 3, center and right panels), consistent with our previous finding that PEG-lipid incorporation suppresses the formation of larger CLs and stabilizes small unilamellar vesicles and bicelles.^57^ The few remaining visible structures in samples prepared with 10 mol% PEG5K-lipid were spherical, while those observed for CLs containing 5 mol% PEG5K-lipid exhibited a more elongated morphology.

**Figure 3.**
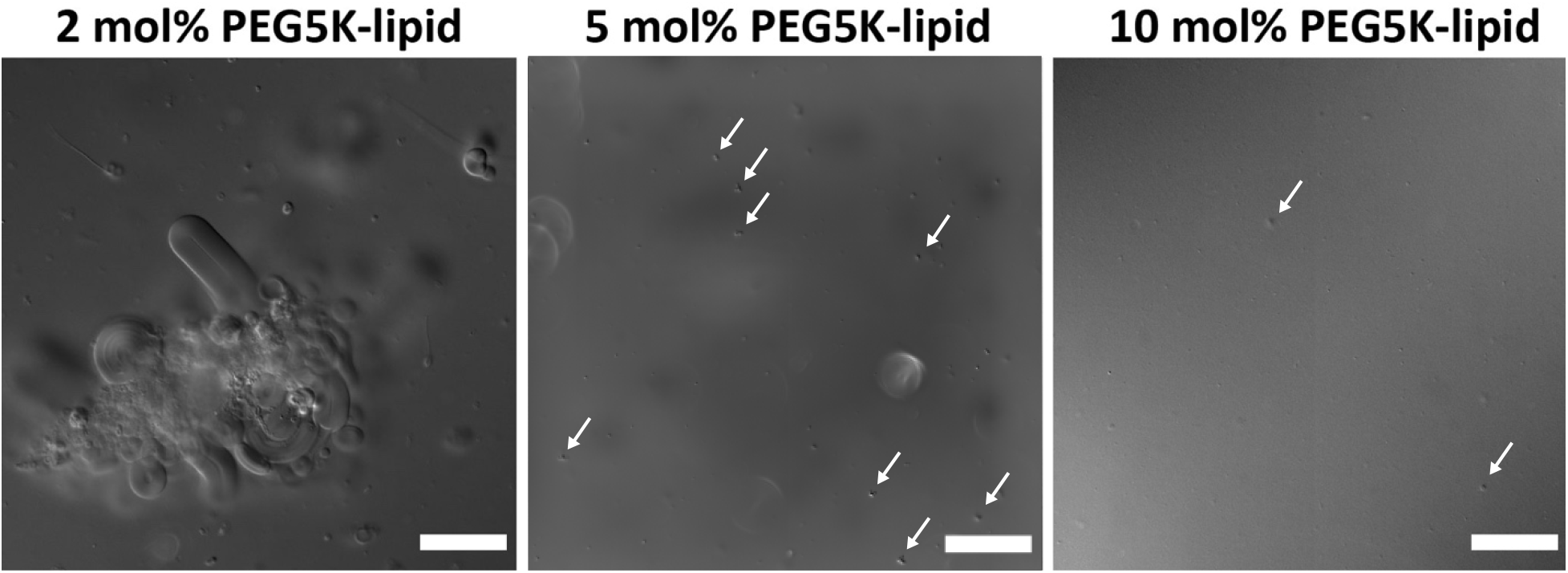
Solubility of highly concentrated lipid formulations as a function of PEG5K-lipid content. DIC microscopy images of sonicated liposomes containing 2, 5, and 10 mol% PEG5K-lipid taken 2 h after rehydration at 67 mM in PBS. Few aggregates or particles greater than ~5 μm (marked by arrows) were observed in liposomes containing 5 and 10 mol% PEG5K-lipid (center and right panels), with the 10 mol% PEG5K-lipid formulation showing the least number of aggregates (i.e., providing strong steric stabilization of lipid particles). In contrast, many aggregates greater than ~5 μm and some greater than 50 μm were present in CLs containing 2 mol% PEG5K-lipid (left panel). CL molar composition: PEG5K-lipid, DOTAP, DOPC, PTX (x_PEG5K-lipid_:50:48-x_PEG5K-lipid_:2). Scale bars: 50 μm.

### Increased PEGylation extends the blood half-life of PTX-loaded fluid-phase CL vectors

Generation of stealth nanoparticles with a prolonged circulation half-life is an important prerequisite for enhanced tumor targeting. To assess the potential of PEGylated CL vectors of PTX for *in vivo* applications, we first studied the effect of differential PEGylation (at 2, 5, or 10 mol% of PEG5K-lipid) on the blood clearance of CL vectors following i.v. administration in immunocompetent healthy mice. The molar composition of the formulations was PEG5K-lipid:DOTAP:DOPC:PTX=x_PEG5K-lipid_:50:48–x_PEG5K-lipid_:2. Compared to the plasma half-lives of the formulations containing 2 and 5 mol% of PEG5K-lipid (39 and 34 min respectively), the half-life of the formulation with 10%of PEG-lipid was extended to 90 min (Figure 4). This increase in half-life was likely due to escape from the surveillance by the reticuloendothelial system, because of an increase in the “stealth” properties of the particles and/or the modulation of the CL shape. Nanoparticle uptake by phagocytic cells is known to be affected by the shape of the particles, a phenomenon attributed to shape- and aspect ratio-dependent activation of different endocytic pathways.^77^

**Figure 4.**
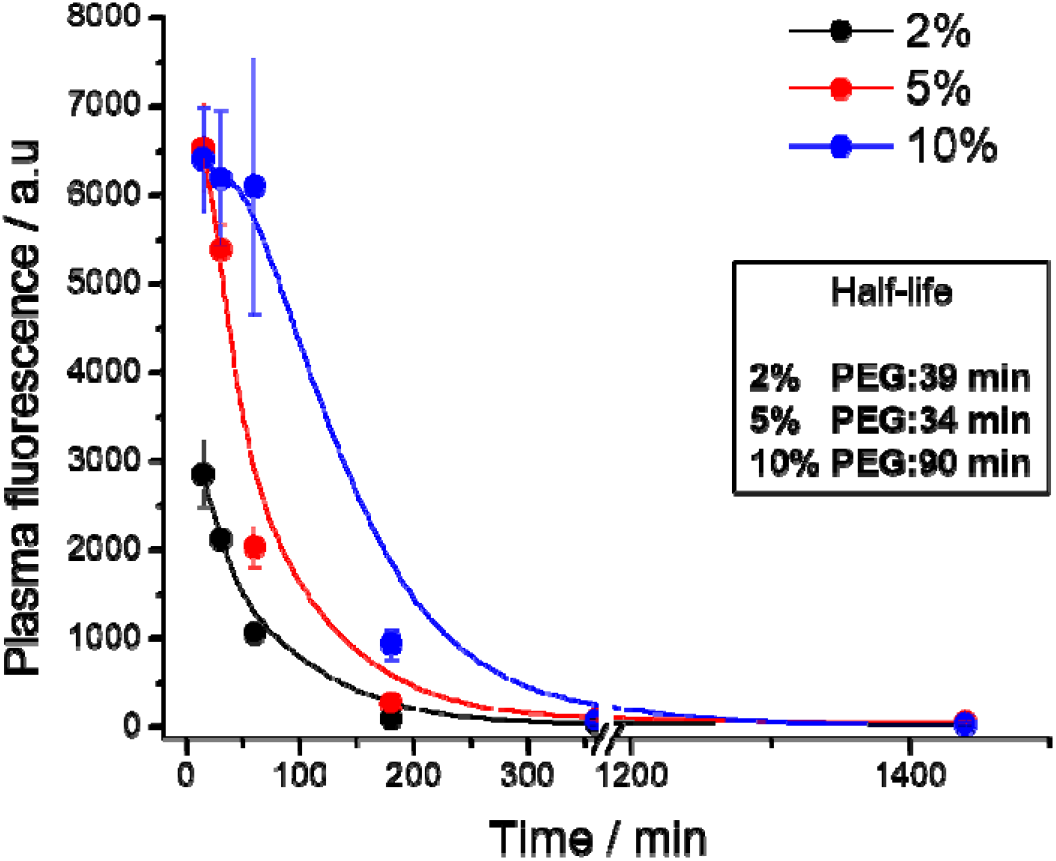
The plasma half-life of PTX-loaded CLs increases with PEGylation. Plasma half-life of i.v. administered FAM-labeled PEGylated PTX-loaded cationic liposome vectors containing 2, 5, and 10 mol% of PEG5K-lipid in healthy immunocompetent BALB/c mice. Mice (*n*=3) were i.v. injected with indicated formulations, blood samples were collected at different time points, and the fluorescence of the plasma was measured. The data were fitted using a biexponential decay formula to obtain the half-life. Error bars indicate the standard error of the mean (±SEM).

The formulation with 10 mol% PEG5K-lipid, with the conformation of the PEG chains far in the brush regime, consists of a mixture of small CLs_PTX_ and in particular fluid-phase nanodiscs_PTX_. The formulation at 5 mol% PEG5K-lipid also consists of a mixture of PEGylated CLs_PTX_ and fluid-phase nanodiscs_PTX_. However, the PEG chains in this case are in the mushroom conformation near the transition to the brush conformation and offer lower levels of steric stabilization compared to the 10 mol% PEG5K-lipid formulation. The 2 mol% PEG5K-lipid formulation, with PEG chains in the mushroom regime far from the brush regime, is not sterically stabilized (consistent with the DIC images in Figure 3).

### Increased PEGylation potentiates tumor accumulation and proapoptotic activity

We next studied the effects of differential PEGylation of PTX-loaded CLs on biodistribution and tumor accumulation. Mice bearing 50–75 mm^3^ orthotopic 4T1 TNBC lesions were i.v. injected with formulations of FAM-labeled CL vectors of PTX at a PEG5K-lipid content of 2, 5, and 10 mol%. Twenty-four hours later, the tumors were collected, cryosectioned, and imaged by confocal fluorescence microscopy (Figure 5A and Figures S5–S14 in the Supporting Information). We observed that the FAM signal in the tumors correlated with the PEG5K-lipid density of the CLs. Specifically, the tumor accumulation of the lipid formulations containing 10 mol% of PEG5K-lipid (PEGylated CLs_PTX_ and coexisting nanodiscsPTX, with the brush conformation of PEG conferring enhanced steric stabilization), was ~10- and ~35-fold higher than that of the formulations containing 5 and 2 mol% of PEG5K-lipid, respectively (Figure 5B).

**Figure 5.**
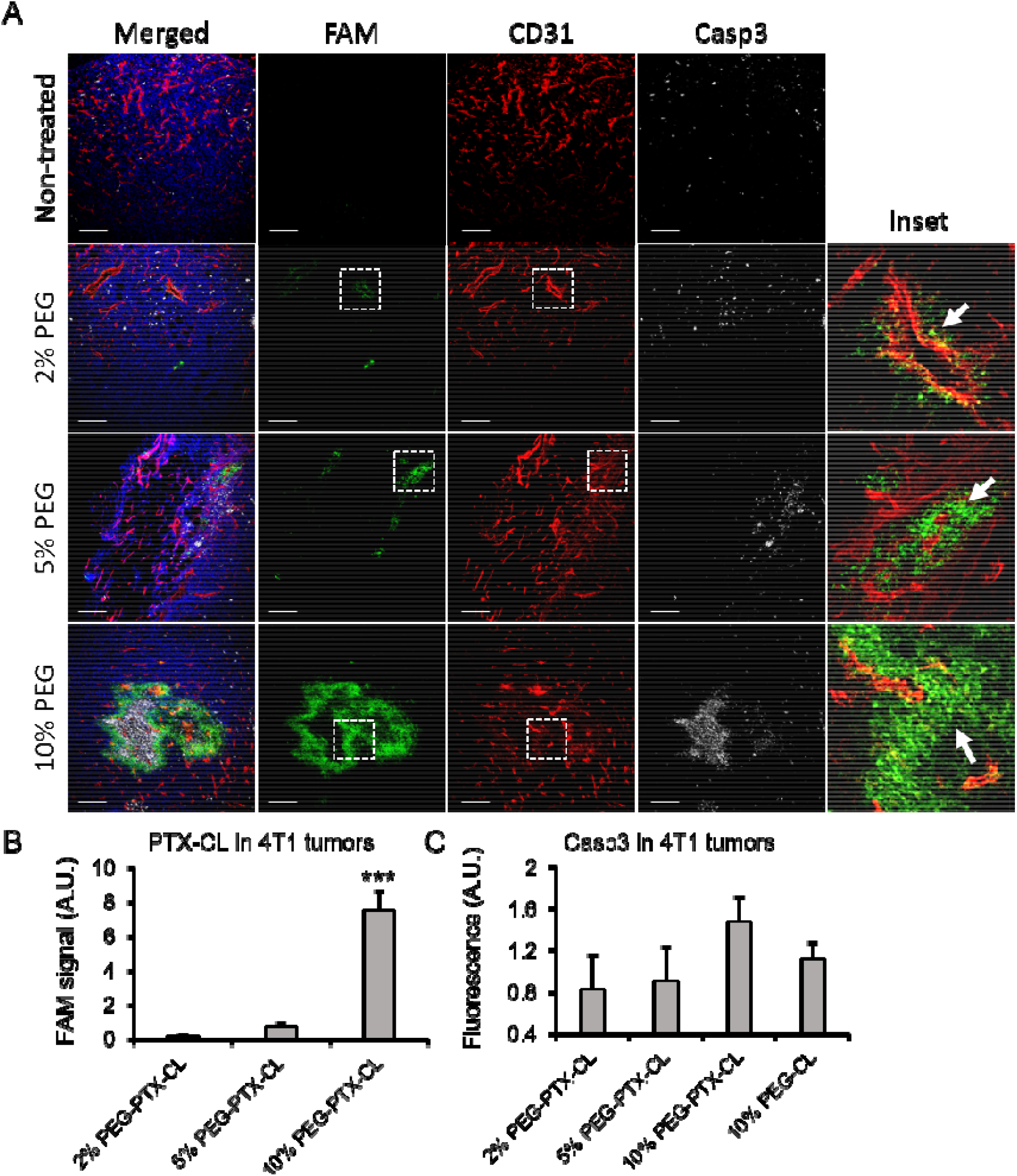
Increased density of PEG5K-lipid in PTX-loaded CLs potentiates accumulation in breast tumors and increases apoptotic cell death. **A:** Representative images of tumor homing and caspase-3 activation by PEGylated CL_PTX_ vectors containing 2, 5, and 10 mol% PEG5K-lipid assessed by immunofluorescence microscopy. The different formulations were administered i.v. in 4T1 tumor-bearing mice; 24 h later the mice were perfused with PBS, tumors excised, cryosectioned, and immunostained for CD31 (blood vessels; red) and cleaved caspase-3 (white), and stained with DAPI nuclear counterstain (blue); the green signal represents the FAM fluorescence of the PEGylated CL vectors of PTX. The FAM signal (lipid vector) observed in the tumors from mice injected with the formulation containing 10 mol% PEG5K-lipid is stronger than that for the formulations with 2 and 5 mol% PEG5K-lipid. In addition, this signal appears to be further from the blood vessels, indicating higher extravasation (white arrows). Scale bars: 200 μm. **B and C:** Quantification of FAM **(B)** and cleaved caspase-3 **(C)** signal, normalized to nontreated tumors from panel A. At least 3 different areas per tumor from 3 different planes and from 3 tumors per groups were analyzed. Error bars indicate the SEM, statistical test: ANOVA one way, Fisher LSD.

The uptake and distribution of drugs and nanoparticles is profoundly affected by the tumor architecture and microenvironment, in particular by the structure and functional status of the tumor vasculature. We next analyzed the fluorescence of the CL vectors in the context of tumor vascular tree (visualized by staining of blood vessels with anti-CD31 antibody) and cellular density (visualized by staining of nuclei with DAPI). We observed that the ability of CL formulations to exit blood vessels and to penetrate into tumor parenchyma was potentiated by the degree of PEGylation. Lipid formulations containing 10 mol% of PEG5K-lipid showed the most extensive extravasation and accumulation of CLs in the tumor parenchyma (Figure 5A insets and Figures S5–S14 in the Supporting Information; white arrows point to the signal from CLs outside the blood vessels). Tumor-driven vascular permeability, commonly seen in rapidly growing solid tumors (including in the TNBC model used in this work),^78,79^ accompanies tumor progression and impacts drug delivery. The small CLsPTX and in particular the fluid-phase nanodiscsPTX in the formulation with 10 mol% PEG5K-lipid, because of their small size and discoidal shape, are expected to have easier access through fenestrated tumor blood vessels than the larger particles in formulations with lower PEGylation density. This, in combination with the extended circulation half-life at 10 mol% PEG5K-lipid, likely accounts for the observed higher extravasation and parenchymal accumulation in the tumor tissue at 10 mol% PEG5K-lipid.

Next, we studied if the increased tumor accumulation of the lipid vector formulation with 10 mol% PEG5K-lipid translated into potentiated *in vivo* anticancer activity. PTX is a mitotic poison that binds to the β subunit of the tubulin heterodimer to prevent depolymerization of microtubules. In 4T1 TNBC cells, the levels of PTX-induced caspase-3 enzymatic activity correlate with tumor sensitivity to the drug and serve as indicator for susceptibility to PTX.^80^ TNBC tumors from mice treated with the PEGylated lipid vectors loaded with PTX were stained for cleaved caspase-3. Whereas a baseline signal of cleaved caspase-3 was observed in the tumors treated with the lipid formulations containing 2 and 5 mol% PEG5K-lipid (Figure 5, panels A and C, and Figures S15-S23 in the Supporting Information), the tumors treated with lipid formulations containing 10 mol% of PEG5K-lipid showed a ~2-fold upregulation of cleaved caspase-3 immunoreactivity (Figure 5C). This difference did not reach statistical significance, possibly due to the low single PTX dose administered and the short follow-up time point. The apoptotic effect in the tumors treated with lipid formulations containing 10 mol% of PEG5K-lipid was due to the PTX, as particles without PTX did not show an increased signal for cleaved caspase-3 (Figure 5C and Figures S24-S26 in the Supporting Information). Further corroborating the ability of the 10 mol% of PEG5K-lipid to efficiently release PTX and trigger cell apoptosis was our observation that caspase-3 was detected in the areas overlapping or adjacent to the fluorescent signal of the lipid formulation (Figure 5A, 4^th^ row, and Figures S15-S23 in the Supporting Information).

In summary, we showed that increasing the content of PEG5K-lipid in CL-based vectors of PTX to the degree that the PEG chains are in the brush conformation not only increases the solubility of the formulation but also the blood half-life, tumor accumulation and tumor penetration of the vector and the anticancer activity of the loaded PTX. The formulation containing CLs with 10 mol% PEG5K-lipid may also be suitable for increasing the solubility of other hydrophobic drugs and potentially find use as a therapeutic carrier for the treatment of solid tumors. To this end, efforts to improve the solubility of PTX in the membranes of CL-based formulations with a high degree of PEGylation are underway. The findings reported here are expected to have far-reaching implications for drug delivery *in vivo* where size, shape, and size stability are critical factors for therapeutic efficacy. Furthermore, a fraction of the pendant PEG moieties may be modified with targeting ligands to achieve active cell-specific targeting.

## Supporting Information

Uncropped versions of the cryo-TEM micrographs in Figure 1 (Figures S1–S4) and confocal microscopy images of stained tumor slices used to obtain the quantification data shown in Figure 5 (Figures S5–S27).

## Acknowledgements

This research study was supported by the National Institutes of Health under award R01GM130769 (CRS, KKE, WF; mechanistic studies on developing lipid nanoparticles for drug delivery), the European Regional Development Fund (TT, Project No. 2014-2020.4.01.15-0012), the Estonian Research Council (TT, grants PRG230 and EAG79; PS, grant PSG38; LSG, grant MOBJD11), EuronanomedII projects ECM-CART and iNanoGun (TT), H2020 MSCA-RISE project Oxigenated (TT), and the Spanish Ministry of Science and Innovation grants RYC2020-028754-I and PID2021-122364OA-I00 (PS). Partial support was provided by the US National Science Foundation (NSF) under Award DMR-1807327 (CRS; kinetic phase behavior of cationic vesicles with incorporated hydrophobic molecules).

## References

(1) Wani, M. C.; Taylor, H. L.; Wall, M. E.; Coggon, P.; McPhail, A. T. Plant antitumor agents. VI. Isolation and structure of taxol, a novel antileukemic and antitumor agent from Taxus brevifolia. J. Am. Chem. Soc. 1971, 93, 2325–2327.

(2) Jordan, M. A.; Wilson, L. Microtubules as a target for anticancer drugs. Nat. Rev. Cancer 2004, 4, 253–265.

(3) Weaver, B. A. How Taxol/paclitaxel kills cancer cells. Mol. Biol. Cell 2014, 25, 2677–2681.

(4) Rowinsky, E. K.; Donehower, R. C. Paclitaxel (Taxol). N. Engl. J. Med. 1995, 332, 1004–1014.

(5) Sofias, A. M.; Dunne, M.; Storm, G.; Allen, C. The battle of “nano” paclitaxel. Adv. Drug Delivery Rev. 2017, 122, 20–30.

(6) Markman, M.; Mekhail, T. M. Paclitaxel in cancer therapy. Expert Opin. Pharmacother. 2002, 3, 755–766.

(7) Ramalingam, S.; Belani, C. P. Paclitaxel for non-small cell lung cancer. Expert Opin. Pharmacother. 2004, 5, 1771–1780.

(8) Hironaka, S.; Zenda, S.; Boku, N.; Fukutomi, A.; Yoshino, T.; Onozawa, Y. Weekly paclitaxel as second-line chemotherapy for advanced or recurrent gastric cancer. Gastric Cancer 2006, 9, 14–18.

(9) Sakamoto, J.; Matsui, T.; Kodera, Y. Paclitaxel chemotherapy for the treatment of gastric cancer. Gastric Cancer 2009, 12, 69–78.

(10) Moxley, K. M.; McMeekin, D. S. Endometrial Carcinoma: A Review of Chemotherapy, Drug Resistance, and the Search for New Agents. Oncologist 2010, 15, 1026–1033.

(11) World Health Organization WHO Model Lists of Essential Medicines, 22nd list (2021). http://who.int/medicines/publications/essentialmedicines/en/ (accessed Jun 16, 2022),

(12) Dranitsaris, G.; Yu, B.; Wang, L.; Sun, W.; Zhou, Y.; King, J.; Kaura, S.; Zhang, A.; Yuan, P. Abraxane^®^ versus Taxol^®^ for patients with advanced breast cancer: A prospective time and motion analysis from a Chinese health care perspective. J. Oncol. Pharm. Pract. 2016, 22, 205–211.

(13) Rugo, H. S.; Barry, W. T.; Moreno-Aspitia, A.; Lyss, A. P.; Cirrincione, C.; Leung, E.; Mayer, E. L.; Naughton, M.; Toppmeyer, D.; Carey, L. A.; Perez, E. A.; Hudis, C.; Winer, E. P. Randomized Phase III Trial of Paclitaxel Once Per Week Compared With Nanoparticle Albumin-Bound Nab-Paclitaxel Once Per Week or Ixabepilone With Bevacizumab As First-Line Chemotherapy for Locally Recurrent or Metastatic Breast Cancer: CALGB 40502/NCCTG N063H (Alliance). J. Clin. Oncol. 2015, 33, 2361–2369.

(14) Gradishar, W. J.; Tjulandin, S.; Davidson, N.; Shaw, H.; Desai, N.; Bhar, P.; Hawkins, M.; O’Shaughnessy, J. Phase III Trial of Nanoparticle Albumin-Bound Paclitaxel Compared With Polyethylated Castor Oil–Based Paclitaxel in Women With Breast Cancer. J. Clin. Oncol. 2005, 23, 7794–7803.

(15) Weiss, R. B.; Donehower, R. C.; Wiernik, P. H.; Ohnuma, T.; Gralla, R. J.; Trump, D. L.; Jr, J. R. B.; Echo, D. A. V.; Hoff, D. D. V.; Leyland-Jones, B. Hypersensitivity reactions from taxol. J. Clin. Oncol. 1990, 8, 1263–1268.

(16) Dorr, R. T. Pharmacology and Toxicology of Cremophor EL Diluent. Ann. Pharmacother. 1994, 28, S11–S14.

(17) Gelderblom, H.; Verweij, J.; Nooter, K.; Sparreboom, A. Cremophor EL: the drawbacks and advantages of vehicle selection for drug formulation. Eur. J. Cancer 2001, 37, 1590–1598.

(18) Hong, S.-S.; Choi, J. Y.; Kim, J. O.; Lee, M.-K.; Kim, S. H.; Lim, S.-J. Development of paclitaxel-loaded liposomal nanocarrier stabilized by triglyceride incorporation. Int. J. Nanomed. 2016, 11, 4465–4477.

(19) Fetterly, G. J.; Straubinger, R. M. Pharmacokinetics of paclitaxel-containing liposomes in rats. AAPS PharmSci 2003, 5, E32.

(20) Sharma, A.; Mayhew, E.; Straubinger, R. M. Antitumor Effect of Taxol-containing Liposomes in a Taxol-resistant Murine Tumor Model. Cancer Res. 1993, 53, 5877–5881.

(21) Zhou, R.; Mazurchuk, R. V.; Tamburlin, J. H.; Harrold, J. M.; Mager, D. E.; Straubinger, R. M. Differential Pharmacodynamic Effects of Paclitaxel Formulations in an Intracranial Rat Brain Tumor Model. J. Pharmacol. Exp. Ther. 2010, 332, 479–488.

(22) Xu, X.; Wang, L.; Xu, H.-Q.; Huang, X.-E.; Qian, Y.-D.; Xiang, J. Clinical comparison between paclitaxel liposome (Lipusu^®^) and paclitaxel for treatment of patients with metastatic gastric cancer. Asian Pac. J. Cancer Prev. 2013, 14, 2591–2594.

(23) Wang, H.; Cheng, G.; Du, Y.; Ye, L.; Chen, W.; Zhang, L.; Wang, T.; Tian, J.; Fu, F. Hypersensitivity reaction studies of a polyethoxylated castor oil-free, liposome-based alternative paclitaxel formulation. Mol. Med. Rep. 2013, 7, 947–952.

(24) Zhang, J. A.; Anyarambhatla, G.; Ma, L.; Ugwu, S.; Xuan, T.; Sardone, T.; Ahmad, I. Development and characterization of a novel Cremophor^®^ EL free liposome-based paclitaxel (LEP-ETU) formulation. Eur. J. Pharm. Biopharm. 2005, 59, 177–187.

(25) Sharma, A.; Sharma, U. S.; Straubinger, R. M. Paclitaxel-liposomes for intracavitary therapy of intraperitoneal P388 leukemia. Cancer Lett. 1996, 107, 265–272.

(26) Simón-Gracia, L.; Hunt, H.; Scodeller, P. D.; Gaitzsch, J.; Braun, G. B.; Willmore, A.-M. A.; Ruoslahti, E.; Battaglia, G.; Teesalu, T. Paclitaxel-Loaded Polymersomes for Enhanced Intraperitoneal Chemotherapy. Mol. Cancer Ther. 2016, 15, 670–679.

(27) Simón-Gracia, L.; Hunt, H.; Scodeller, P.; Gaitzsch, J.; Kotamraju, V. R.; Sugahara, K. N.; Tammik, O.; Ruoslahti, E.; Battaglia, G.; Teesalu, T. iRGD peptide conjugation potentiates intraperitoneal tumor delivery of paclitaxel with polymersomes. Biomaterials 2016, 104, 247–257.

(28) Safinya, C. R.; Ewert, K. K.; Majzoub, R. N.; Leal, C. Cationic liposome-nucleic acid complexes for gene delivery and gene silencing. New J. Chem. 2014, 38, 5164–5172.

(29) Ewert, K. K.; Scodeller, P.; Simón-Gracia, L.; Steffes, V. M.; Wonder, E. A.; Teesalu, T.; Safinya, C. R. Cationic Liposomes as Vectors for Nucleic Acid and Hydrophobic Drug Therapeutics. Pharmaceutics 2021, 13, 1365.

(30) Torchilin, V. P. Recent advances with liposomes as pharmaceutical carriers. Nat. Rev. Drug Discovery 2005, 4, 145.

(31) Allen, T. M.; Cullis, P. R. Liposomal drug delivery systems: From concept to clinical applications. Adv. Drug Delivery Rev. 2013, 65, 36–48.

(32) Witzigmann, D.; Kulkarni, J. A.; Leung, J.; Chen, S.; Cullis, P. R.; van der Meel, R. Lipid nanoparticle technology for therapeutic gene regulation in the liver. Adv. Drug Delivery Rev. 2020, 159, 344–363.

(33) Kulkarni, J. A.; Witzigmann, D.; Chen, S.; Cullis, P. R.; van der Meel, R. Lipid Nanoparticle Technology for Clinical Translation of siRNA Therapeutics. Acc. Chem. Res. 2019, 52, 2435–2444.

(34) Sercombe, L.; Veerati, T.; Moheimani, F.; Wu, S. Y.; Sood, A. K.; Hua, S. Advances and Challenges of Liposome Assisted Drug Delivery. Front. Pharmacol. 2015, 6, 286.

(35) Koudelka, Š.; Turánek, J. Liposomal paclitaxel formulations. J. Controlled Release 2012, 163, 322–334.

(36) Campbell, R. B.; Ying, B.; Kuesters, G. M.; Hemphill, R. Fighting cancer: From the bench to bedside using second generation cationic liposomal therapeutics. J. Pharm. Sci. 2009, 98, 411–429.

(37) Barenholz, Y. Doxil^®^— The first FDA-approved nano-drug: Lessons learned. J. Controlled Release 2012, 160, 117–134.

(38) Lasic, D. D.; Čeh, B.; Stuart, M. C. A.; Guo, L.; Frederik, P. M.; Barenholz, Y. Transmembrane gradient driven phase transitions within vesicles: lessons for drug delivery. Biochim. Biophys. Acta, Biomembr. 1995, 1239, 145–156.

(39) Mulligan, M. J.; Lyke, K. E.; Kitchin, N.; Absalon, J.; Gurtman, A.; Lockhart, S.; Neuzil, K.; Raabe, V.; Bailey, R.; Swanson, K. A.; Li, P.; Koury, K.; Kalina, W.; Cooper, D.; Fontes-Garfias, C.; Shi, P.-Y.; Türeci, Ö.; Tompkins, K. R.; Walsh, E. E.; Frenck, R.; Falsey, A. R.; Dormitzer, P. R.; Gruber, W. C.; Şahin, U.; Jansen, K. U. Phase I/II study of COVID-19 RNA vaccine BNT162b1 in adults. Nature 2020, 586, 589–593.

(40) Polack, F. P.; Thomas, S. J.; Kitchin, N.; Absalon, J.; Gurtman, A.; Lockhart, S.; Perez, J. L.; Pérez Marc, G.; Moreira, E. D.; Zerbini, C.; Bailey, R.; Swanson, K. A.; Roychoudhury, S.; Koury, K.; Li, P.; Kalina, W. V.; Cooper, D.; Frenck, R. W.; Hammitt, L. L.; Türeci, Ö.; Nell, H.; Schaefer, A.;Ünal, S.; Tresnan, D. B.; Mather, S.; Dormitzer, P. R.; Şahin, U.; Jansen, K. U.; Gruber, W. C. Safety and Efficacy of the BNT162b2 mRNA Covid-19 Vaccine. N. Engl. J. Med. 2020, 383, 2603–2615.

(41) Jackson, L. A.; Anderson, E. J.; Rouphael, N. G.; Roberts, P. C.; Makhene, M.; Coler, R. N.; McCullough, M. P.; Chappell, J. D.; Denison, M. R.; Stevens, L. J.; Pruijssers, A. J.; McDermott, A.; Flach, B.; Doria-Rose, N. A.; Corbett, K. S.; Morabito, K. M.; O’Dell, S.; Schmidt, S. D.; Swanson, P. A.; Padilla, M.; Mascola, J. R.; Neuzil, K. M.; Bennett, H.; Sun, W.; Peters, E.; Makowski, M.; Albert, J.; Cross, K.; Buchanan, W.; Pikaart-Tautges, R.; Ledgerwood, J. E.; Graham, B. S.; Beigel, J. H. An mRNA Vaccine against SARS-CoV-2 — Preliminary Report. N. Engl. J. Med. 2020, 383, 1920–1931.

(42) Baden, L. R.; El Sahly, H. M.; Essink, B.; Kotloff, K.; Frey, S.; Novak, R.; Diemert, D.; Spector, S. A.; Rouphael, N.; Creech, C. B.; McGettigan, J.; Khetan, S.; Segall, N.; Solis, J.; Brosz, A.; Fierro, C.; Schwartz, H.; Neuzil, K.; Corey, L.; Gilbert, P.; Janes, H.; Follmann, D.; Marovich, M.; Mascola, J.; Polakowski, L.; Ledgerwood, J.; Graham, B. S.; Bennett, H.; Pajon, R.; Knightly, C.; Leav, B.; Deng, W.; Zhou, H.; Han, S.; Ivarsson, M.; Miller, J.; Zaks, T. Efficacy and Safety of the mRNA-1273 SARS-CoV-2 Vaccine. N. Engl. J. Med. 2020, 384, 403–416.

(43) Fasol, U.; Frost, A.; Büchert, M.; Arends, J.; Fiedler, U.; Scharr, D.; Scheuenpflug, J.; Mross, K. Vascular and pharmacokinetic effects of EndoTAG-1 in patients with advanced cancer and liver metastasis. Ann. Oncol. 2012, 23, 1030–1036.

(44) Strieth, S.; Eichhorn, M. E.; Sauer, B.; Schulze, B.; Teifel, M.; Michaelis, U.; Dellian, M. Neovascular targeting chemotherapy: Encapsulation of paclitaxel in cationic liposomes impairs functional tumor microvasculature. Int. J. Cancer 2004, 110, 117–124.

(45) Strieth, S.; Nussbaum, C. F.; Eichhorn, M. E.; Fuhrmann, M.; Teifel, M.; Michaelis, U.; Berghaus, A.; Dellian, M. Tumor-selective vessel occlusions by platelets after vascular targeting chemotherapy using paclitaxel encapsulated in cationic liposomes. Int. J. Cancer 2008, 122, 452–460.

(46) Kunstfeld, R.; Wickenhauser, G.; Michaelis, U.; Teifel, M.; Umek, W.; Naujoks, K.; Wolff, K.; Petzelbauer, P. Paclitaxel Encapsulated in Cationic Liposomes Diminishes Tumor Angiogenesis and Melanoma Growth in a “Humanized” SCID Mouse Model. J. Invest. Dermatol. 2003, 120, 476–482.

(47) Schmitt-Sody, M.; Strieth, S.; Krasnici, S.; Sauer, B.; Schulze, B.; Teifel, M.; Michaelis, U.; Naujoks, K.; Dellian, M. Neovascular Targeting Therapy: Paclitaxel Encapsulated in Cationic Liposomes Improves Antitumoral Efficacy. Clin. Cancer Res. 2003, 9, 2335–2341.

(48) Zhen, Y.; Ewert, K. K.; Fisher, W. S.; Steffes, V. M.; Li, Y.; Safinya, C. R. Paclitaxel loading in cationic liposome vectors is enhanced by replacement of oleoyl with linoleoyl tails with distinct lipid shapes. Sci. Rep. 2021, 11, 7311.

(49) Ewert, K. K.; Kotamraju, V. R.; Majzoub, R. N.; Steffes, V. M.; Wonder, E. A.; Teesalu, T.; Ruoslahti, E.; Safinya, C. R. Synthesis of linear and cyclic peptide–PEG–lipids for stabilization and targeting of cationic liposome–DNA complexes. Bioorg. Med. Chem. Lett. 2016, 26, 1618–1623.

(50) Wonder, E.; Simón-Gracia, L.; Scodeller, P.; Majzoub, R. N.; Kotamraju, V. R.; Ewert, K. K.; Teesalu, T.; Safinya, C. R. Competition of charge-mediated and specific binding by peptide-tagged cationic liposome–DNA nanoparticles in vitro and in vivo. Biomaterials 2018, 166, 52–63.

(51) Koltover, I.; Salditt, T.; Rädler, J. O.; Safinya, C. R. An inverted hexagonal phase of cationic liposome-DNA complexes related to DNA release and delivery. Science 1998, 281, 78–81.

(52) Ewert, K. K.; Evans, H. M.; Zidovska, A.; Bouxsein, N. F.; Ahmad, A.; Safinya, C. R. A columnar phase of dendritic lipid-based cationic liposome-DNA complexes for gene delivery: Hexagonally ordered cylindrical micelles embedded in a DNA honeycomb lattice. J. Am. Chem. Soc. 2006, 128, 3998–4006.

(53) Leal, C.; Bouxsein, N. F.; Ewert, K. K.; Safinya, C. R. Highly Efficient Gene Silencing Activity of siRNA Embedded in a Nanostructured Gyroid Cubic Lipid Matrix. J. Am. Chem. Soc. 2010, 132, 16841–16847.

(54) Zidovska, A.; Evans, H. M.; Ewert, K. K.; Quispe, J.; Carragher, B.; Potter, C. S.; Safinya, C. R. Liquid crystalline phases of dendritic lipid-DNA self-assemblies: Lamellar, hexagonal, and DNA bundles. J. Phys. Chem. B 2009, 113, 3694–3703.

(55) Safinya, C. R.; Ewert, K. K.; Li, Y.; Rädler, J. O., Cationic Liposomes as Spatial Organizers of Nucleic Acids in One, Two, and Three Dimensions: Liquid Crystal Phases with Applications in Delivery and Bionanotechnology. In Handbook of Lipid Membranes: Molecular, Functional, and Materials Aspects, Safinya, C. R.; Rädler, J. O., Eds. CRC Press, Taylor & Francis Group: Boca Raton, 2021; pp 195–209.

(56) Majzoub, R. N.; Ewert, K. K.; Jacovetty, E. L.; Carragher, B.; Potter, C. S.; Li, Y.; Safinya, C. R. Patterned Threadlike Micelles and DNA-Tethered Nanoparticles: A Structural Study of PEGylated Cationic Liposome–DNA Assemblies. Langmuir 2015, 31, 7073–7083.

(57) Steffes, V. M.; Zhang, Z.; MacDonald, S.; Crowe, J.; Ewert, K. K.; Carragher, B.; Potter, C. S.; Safinya, C. R. PEGylation of Paclitaxel-Loaded Cationic Liposomes Drives Steric Stabilization of Bicelles and Vesicles thereby Enhancing Delivery and Cytotoxicity to Human Cancer Cells. ACS Appl. Mater. Interfaces 2020, 12, 151–162.

(58) Helfrich, W. Z. Elastic Properties of Lipid Bilayers: Theory and Possible Experiments. Z. Naturforsch., C: J. Biosci. 1973, 28, 693–703.

(59) Lipowsky, R.; Sackmann, E., Structure and Dynamics of Membranes. Elsevier: Amsterdam, 1995; Vol. 1A.

(60) Safran, S. A., Statistical thermodynamics of surfaces, interfaces, and membranes. Addison-Wesley: Reading, 1994; Vol. 90.

(61) Israelachvili, J. N.; Mitchell, D. J.; Ninham, B. W. Theory of self-assembly of hydrocarbon amphiphiles into micelles and bilayers. J. Chem. Soc.-Faraday Trans. 2 1976, 72, 1525–1568.

(62) Israelachvili, J. N.; Mitchell, D. J.; Ninham, B. W. Theory of self-assembly of lipid bilayers and vesicles. Biochim. Biophys. Acta, Biomembr. 1977, 470, 185–201.

(63) Israelachvili, J. N., Intermolecular and Surface Forces. 3rd ed.; Elsevier: Amsterdam, 2011.

(64) Johnsson, M.; Edwards, K. Liposomes, Disks, and Spherical Micelles: Aggregate Structure in Mixtures of Gel Phase Phosphatidylcholines and Poly(Ethylene Glycol)-Phospholipids. Biophys. J. 2003, 85, 3839–3847.

(65) Aresh, W.; Liu, Y.; Sine, J.; Thayer, D.; Puri, A.; Huang, Y.; Wang, Y.; Nieh, M.-P. The Morphology of Self-Assembled Lipid-Based Nanoparticles Affects Their Uptake by Cancer Cells. J. Biomed. Nanotechnol. 2016, 12, 1852–1863.

(66) Rad, A. T.; Chen, C.-W.; Aresh, W.; Xia, Y.; Lai, P.-S.; Nieh, M.-P. Combinational Effects of Active Targeting, Shape, and Enhanced Permeability and Retention for Cancer Theranostic Nanocarriers. ACS Appl. Mater. Interfaces 2019, 11, 10505–10519.

(67) Rad, A. T.; Hargrove, D.; Daneshmandi, L.; Ramsdell, A.; Lu, X.; Nieh, M.-P. Codelivery of Paclitaxel and Parthenolide in Discoidal Bicelles for a Synergistic Anticancer Effect: Structure Matters. Adv. NanoBiomed Res. 2022, 2, 2100080.

(68) Smith, G. S.; Sirota, E. B.; Safinya, C. R.; Plano, R. J.; Clark, N. A. X-ray structural studies of freely suspended ordered hydrated DMPC multimembrane films. J. Chem. Phys. 1990, 92, 4519–4529.

(69) Campbell, R. B.; Balasubramanian, S. V.; Straubinger, R. M. Influence of cationic lipids on the stability and membrane properties of paclitaxel-containing liposomes. J. Pharm. Sci. 2001, 90, 1091–1105.

(70) Kannan, V.; Balabathula, P.; Divi, M. K.; Thoma, L. A.; Wood, G. C. Optimization of drug loading to improve physical stability of paclitaxel-loaded long-circulating liposomes. J. Liposome Res. 2015, 25, 308–315.

(71) Bernsdorff, C.; Reszka, R.; Winter, R. Interaction of the anticancer agent Taxol™ (paclitaxel) with phospholipid bilayers. J. Biomed. Mater. Res. 1999, 46, 141–149.

(72) Steffes, V. M.; Murali, M. M.; Park, Y.; Fletcher, B. J.; Ewert, K. K.; Safinya, C. R. Distinct Solubility and Cytotoxicity Regimes of Paclitaxel-Loaded Cationic Liposomes at Low and High Drug Content Revealed by Kinetic Phase Behavior and Cancer Cell Viability Studies. Biomaterials 2017, 145, 242–255.

(73) Balasubramanian, S. V.; Straubinger, R. M. Taxol-Lipid Interactions: Taxol-Dependent Effects on the Physical Properties of Model Membranes. Biochemistry 1994, 33, 8941–8947.

(74) Di, J.; Gao, X.; Du, Y.; Zhang, H.; Gao, J.; Zheng, A. Size, shape, charge and “stealthy” surface: Carrier properties affect the drug circulation time in vivo. Asian J. Pharm. Sci. (Shenyang, China) 2021, 16, 444–458.

(75) Eichhorn, M. E.; Ischenko, I.; Luedemann, S.; Strieth, S.; Papyan, A.; Werner, A.; Bohnenkamp, H.; Guenzi, E.; Preissler, G.; Michaelis, U.; Jauch, K.-W.; Bruns, C. J.; Dellian, M. Vascular targeting by EndoTAG™-1 enhances therapeutic efficacy of conventional chemotherapy in lung and pancreatic cancer. Int. J. Cancer 2010, 126, 1235–1245.

(76) Eichhorn, M. E.; Luedemann, S.; Strieth, S.; Papyan, A.; Ruhstorfer, H.; Haas, H.; Michaelis, U.; Sauer, B.; Teifel, M.; Enders, G. H.; Brix, G.; Jauch, K. W.; Bruns, C. J.; Dellian, M. Cationic lipid complexed camptothecin (EndoTAG^®^-2) improves antitumoral efficacy by tumor vascular targeting. Cancer Biol. Ther. 2007, 6, 920–929.

(77) Li, Z.; Sun, L.; Zhang, Y.; Dove, A. P.; O’Reilly, R. K.; Chen, G. Shape Effect of Glyco-Nanoparticles on Macrophage Cellular Uptake and Immune Response. ACS Macro Lett. 2016, 5, 1059–1064.

(78) Lepland, A.; Malfanti, A.; Haljasorg, U.; Asciutto, E. K.; Pickholz, M.; Bringas, M.; Đorđević, S.; Salumäe, L.; Peterson, P.; Teesalu, T.; Vicent, M. J.; Scodeller, P. Depletion of Mannose Receptor–Positive Tumor-associated Macrophages via a Peptide-targeted Star-shaped Polyglutamate Inhibits Breast Cancer Progression in Mice. Cancer Res. Commun. 2022, 2, 533–551.

(79) Simón-Gracia, L.; Scodeller, P.; Fuentes, S. S.; Gómez, V.; Vallejo, X. R.; San Sebastián, E.; Sidorenko, V.; Di Silvio, D.; Suck, M.; De Lorenzi, F.; Rizzo, L. Y.; von Stillfried, S.; Kilk, K.; Lammers, T.; Moya, S. E.; Teesalu, T. Application of polymersomes engineered to target p32 protein for detection of small breast tumors in mice. Oncotarget 2018, 9, 18682–18697.

(80) Odonkor, C. A.; Achilefu, S. Differential Activity of Caspase-3 Regulates Susceptibility of Lung and Breast Tumor Cell Lines to Paclitaxel. Open Biochem. J. 2008, 2, 121–128.

