## Supporting Information for "Paclitaxel-loaded Cationic Fluid Lipid Nanodiscs and Liposomes with Brush-Conformation PEG Chains Penetrate Breast Tumors and Trigger Caspase-3 Activation"

for

### Contents


| | |
|---|---|
| Uncropped cryogenic TEM images | Figure S1–S4 |
| Confocal microscopy images of stained tumor slices | Figure S5–S27 |

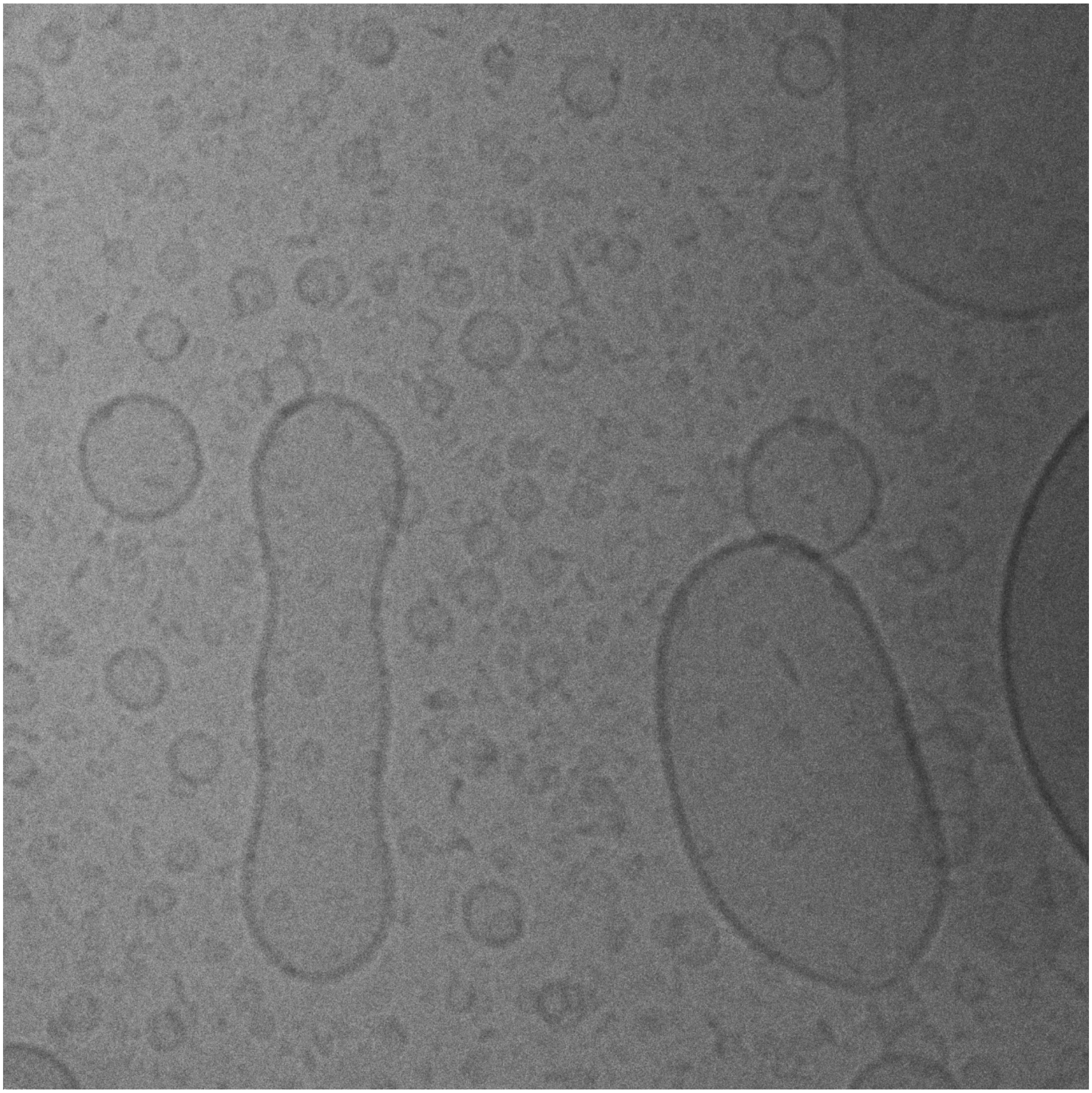

**Figure S1.** Uncropped cryogenic TEM image of PTX-loaded CLs without PEGylation (top left panel in Figure 1). The sonicated liposomes at the EndoTAG-1 composition (DOTAP/DOPC/PTX=50/47/3 molar ratio) exhibit both larger and smaller vesicles with few discs.

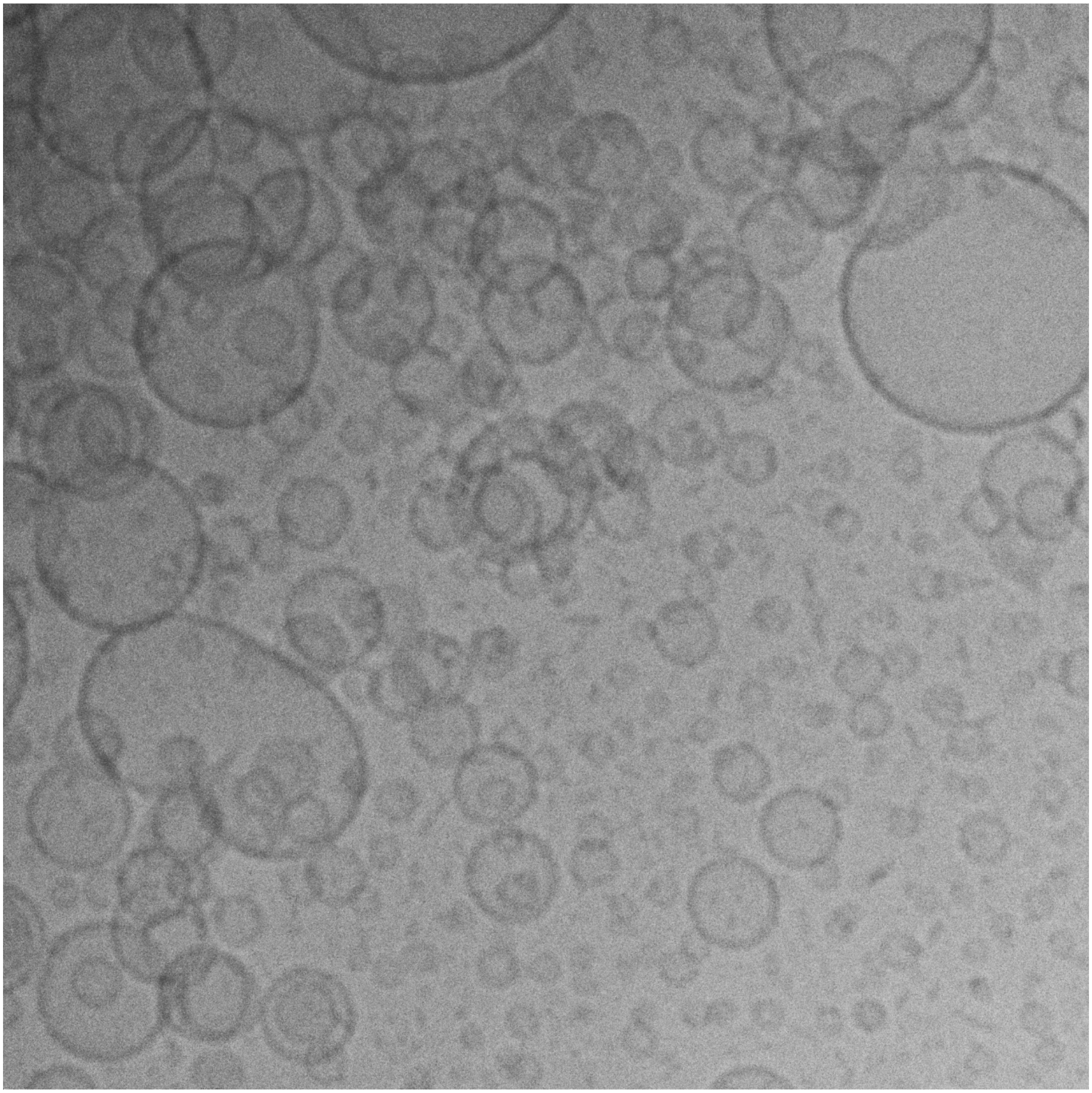

**Figure S2.** Uncropped cryogenic TEM image of PTX-loaded CLs without PEGylation (bottom left panel in Figure 1). The sonicated liposomes at the EndoTAG-1 composition (DOTAP/DOPC/PTX=50/47/3 molar ratio) exhibit both larger and smaller vesicles with few discs.

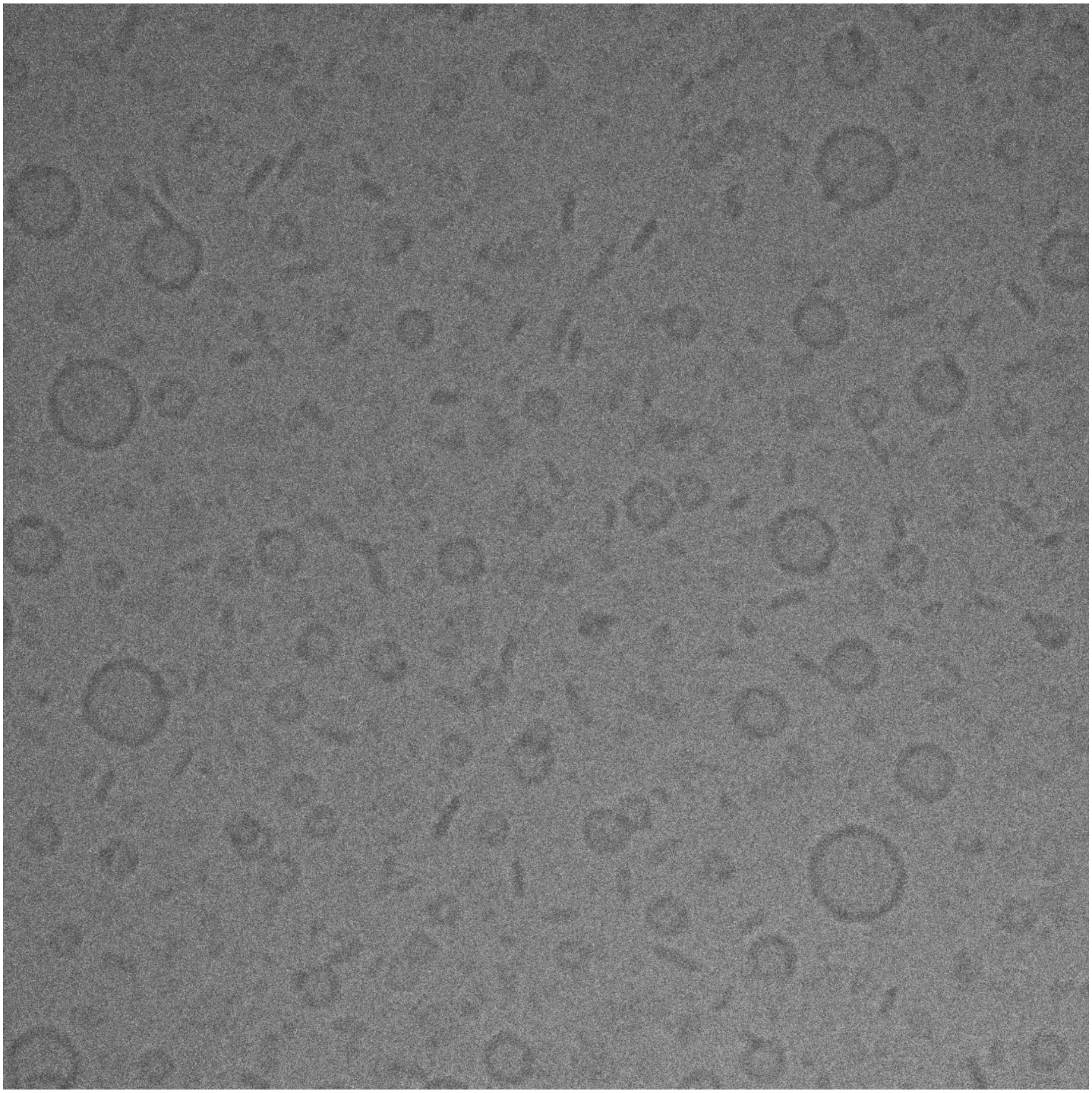

**Figure S3.** Uncropped cryogenic TEM image of PEGylated PTX-loaded CLs (top right panel in Figure 1). The sonicated PTX-loaded CLs with 10 mol% PEG2K-lipid (DOTAP/DOPC/PEG2K-lipid/PTX=50/37/10/3 molar ratio) lack vesicles above ~50 nm in size and show a prevalence of very small vesicles and lipid discs.

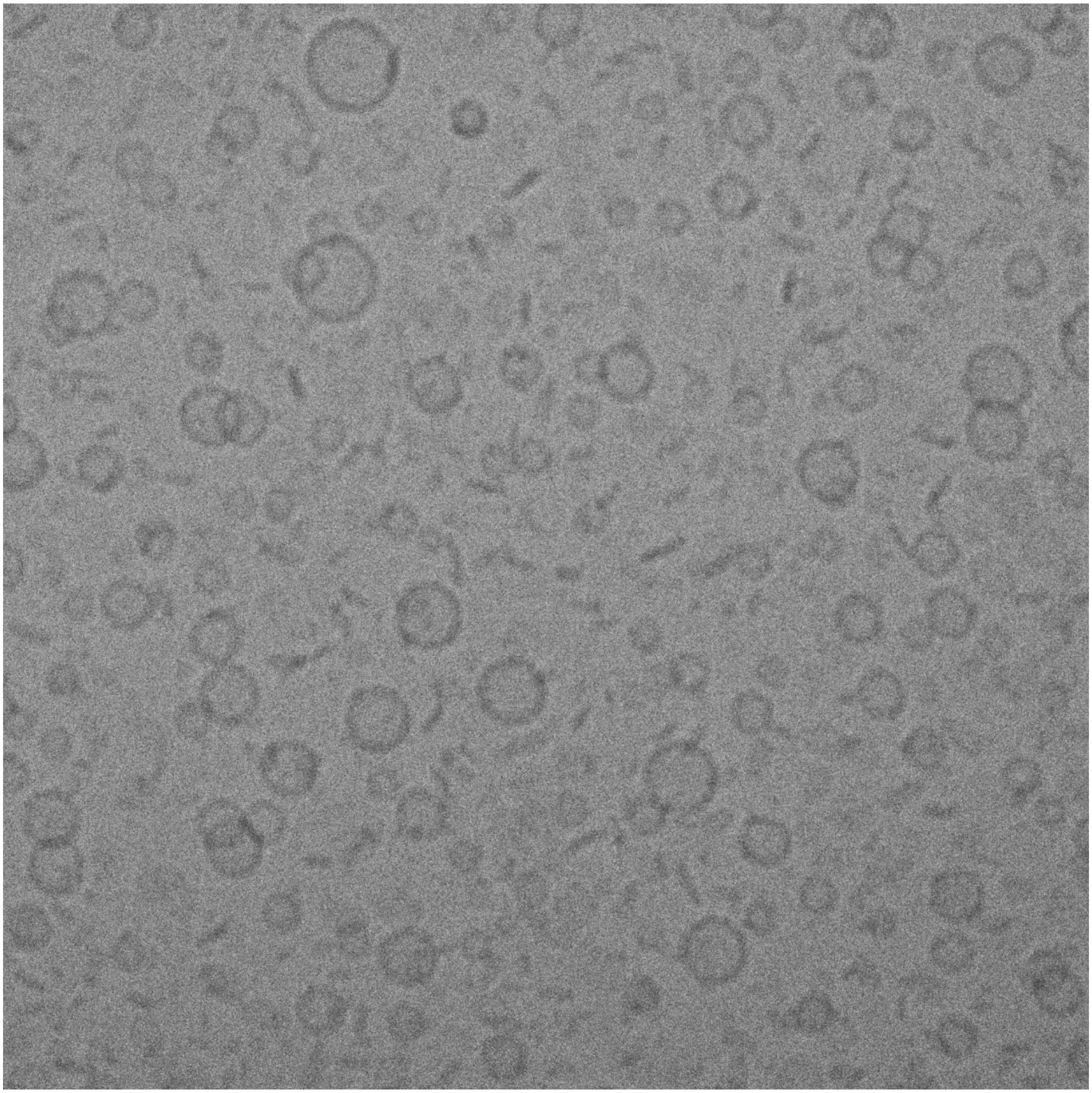

**Figure S4.** Uncropped cryogenic TEM image of PEGylated PTX-loaded CLs (bottom right panel in Figure 1). The sonicated PTX-loaded CLs with 10 mol% PEG2K-lipid (DOTAP/DOPC/PEG2K-lipid/PTX=50/37/10/3 molar ratio) lack vesicles above ~50 nm in size and show a prevalence of very small vesicles and lipid discs.

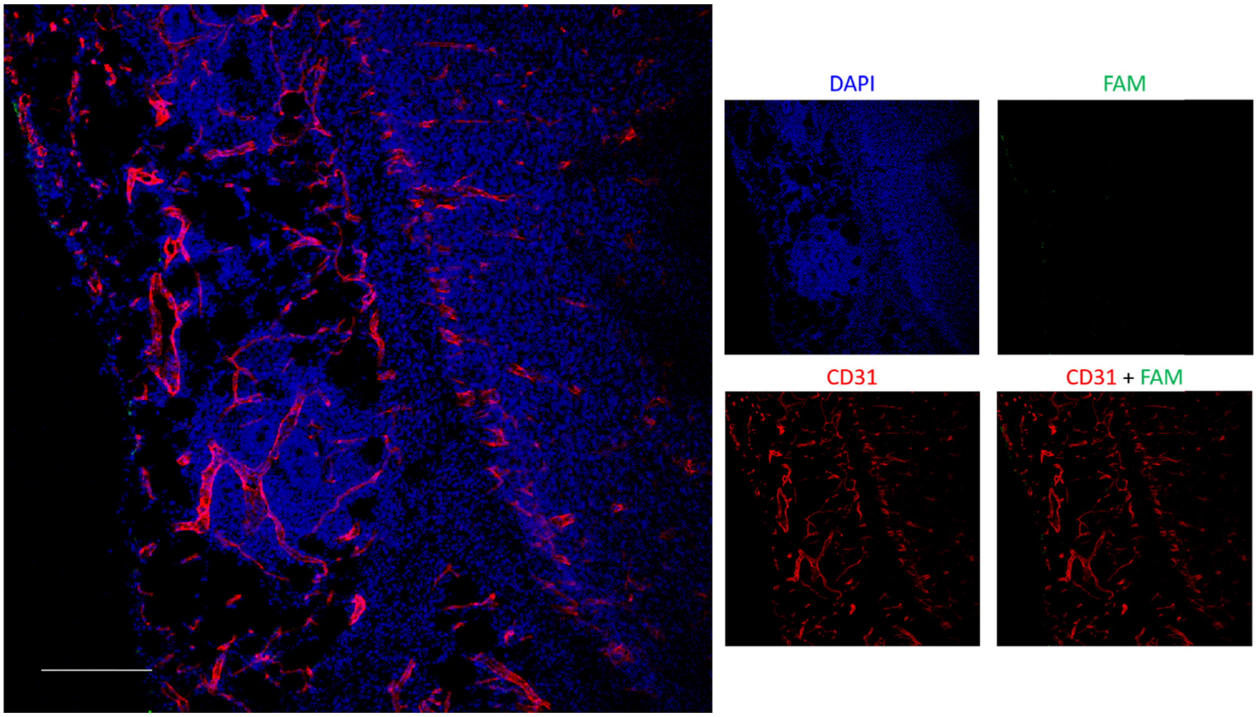


**Figure S5.** Tumor homing of PEGylated $CL_{PTX}$ vector containing 2 mol% PEG5K-lipid assessed by immunofluorescence microscopy. The formulation was administered intravenously in 4T1 tumor-bearing mouse; 24 h later the mouse was perfused with PBS, tumor excised, cryo-sectioned, and immunostained for CD31 (blood vessels; red) and stained with DAPI nuclear counterstain (blue); the green signal represents the FAM fluorescence of the PEG-$CL_{PTX}$. Tumor from mouse #1. Scale bar: 200 µm.

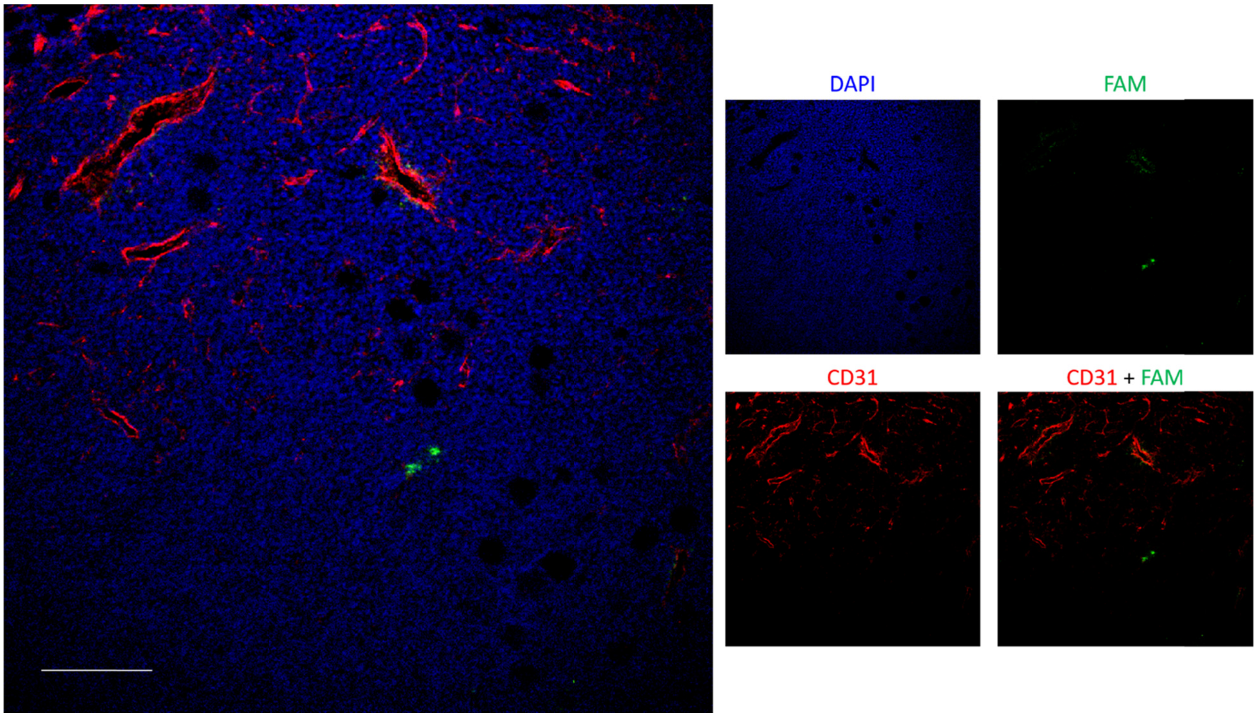


**Figure S6.** Tumor homing of PEGylated $CL_{PTX}$ vector containing 2 mol% PEG5K-lipid assessed by immunofluorescence microscopy. The formulation was administered intravenously in 4T1 tumor-bearing mouse; 24 h later the mouse was perfused with PBS, tumor excised, cryo-sectioned, and immunostained for CD31 (blood vessels; red) and stained with DAPI nuclear counterstain (blue); the green signal represents the FAM fluorescence of the PEG-$CL_{PTX}$. Tumor from mouse #2. Scale bar: 200 µm.

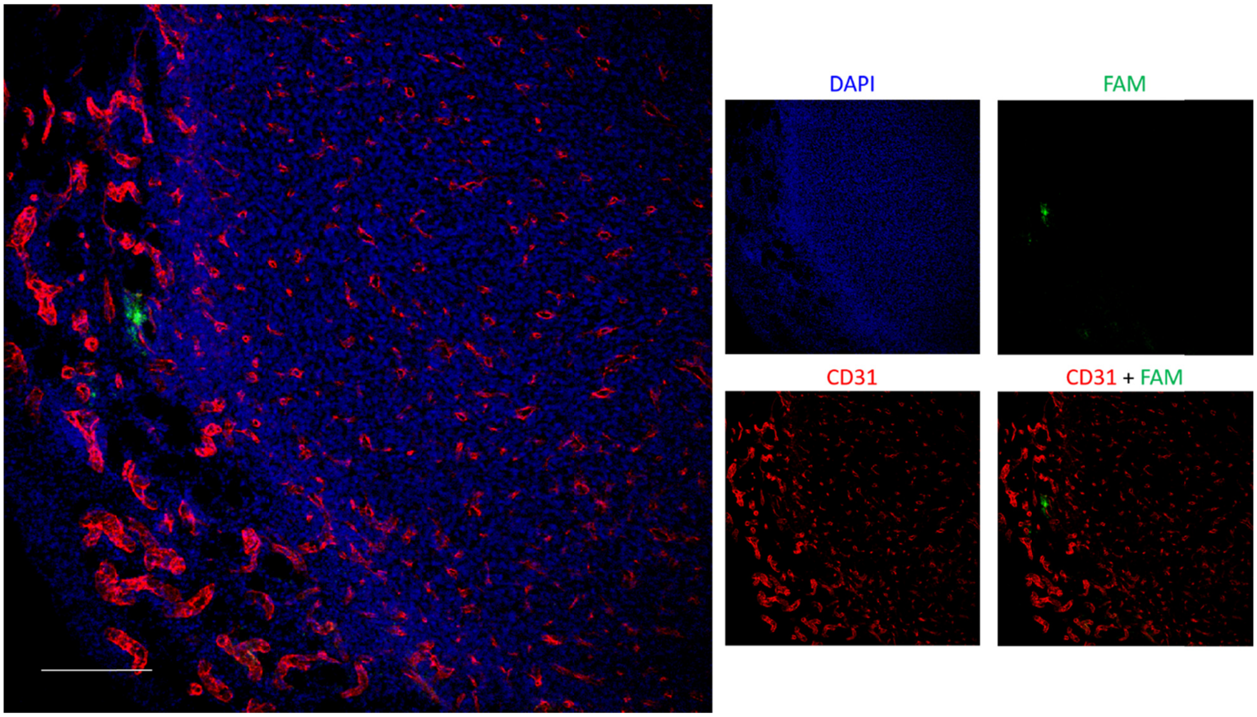


**Figure S7.** Tumor homing of PEGylated $CL_{PTX}$ vector containing 2 mol% PEG5K-lipid assessed by immunofluorescence microscopy. The formulation was administered intravenously in 4T1 tumor-bearing mouse; 24 h later the mouse was perfused with PBS, tumor excised, cryo-sectioned, and immunostained for CD31 (blood vessels; red) and stained with DAPI nuclear counterstain (blue); the green signal represents the FAM fluorescence of the PEG-$CL_{PTX}$. Tumor from mouse #3. Scale bar: 200 µm.

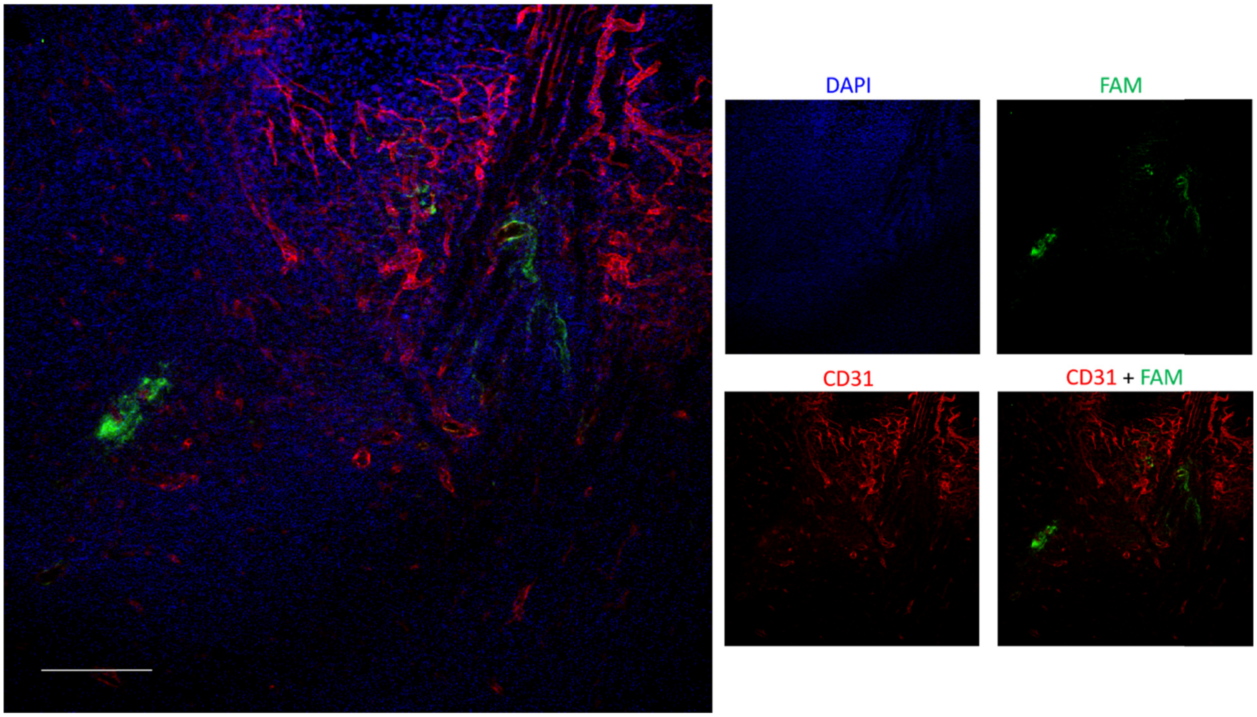


**Figure S8.** Tumor homing of PEGylated $CL_{PTX}$ vector containing 5 mol% PEG5K-lipid assessed by immunofluorescence microscopy. The formulation was administered intravenously in 4T1 tumor-bearing mouse; 24 h later the mouse was perfused with PBS, tumor excised, cryo-sectioned, and immunostained for CD31 (blood vessels; red) and stained with DAPI nuclear counterstain (blue); the green signal represents the FAM fluorescence of the PEG-$CL_{PTX}$. Tumor from mouse #4. Scale bar: 200 µm.

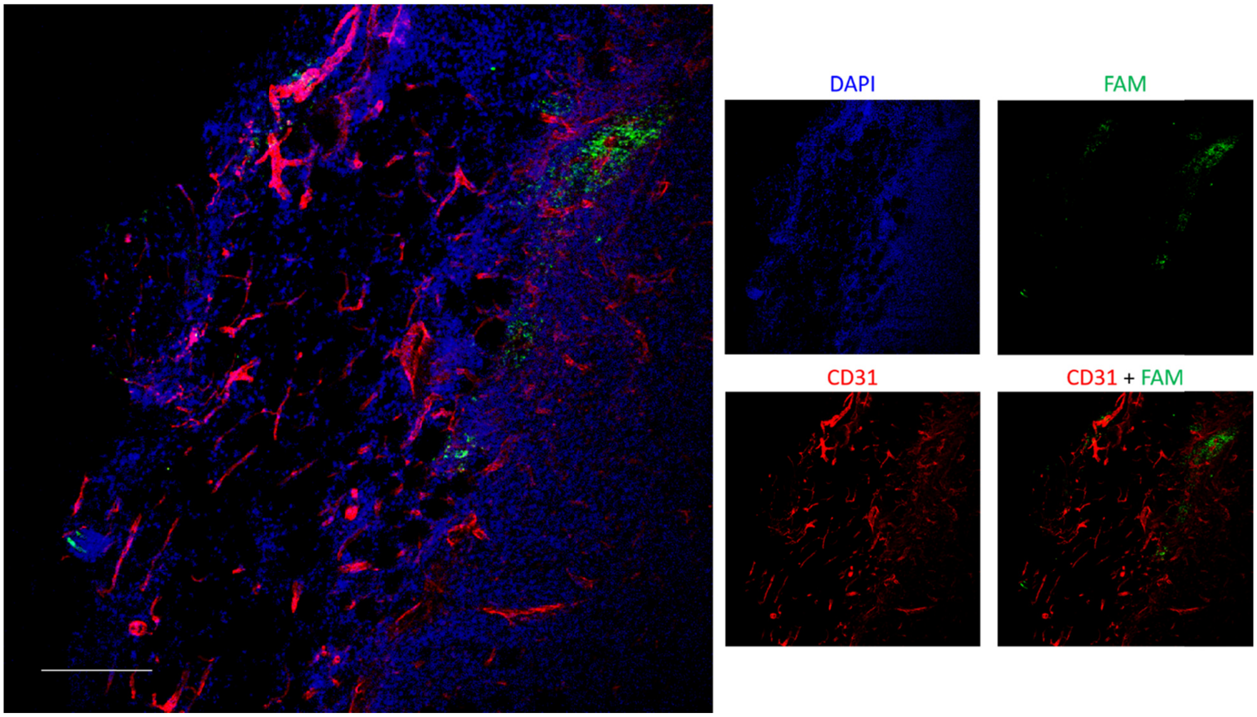


**Figure S9.** Tumor homing of PEGylated $CL_{PTX}$ vector containing 5 mol% PEG5K-lipid assessed by immunofluorescence microscopy. The formulation was administered intravenously in 4T1 tumor-bearing mouse; 24 h later the mouse was perfused with PBS, tumor excised, cryo-sectioned, and immunostained for CD31 (blood vessels; red) and stained with DAPI nuclear counterstain (blue); the green signal represents the FAM fluorescence of the PEG-$CL_{PTX}$. Tumor from mouse #5. Scale bar: 200 µm.

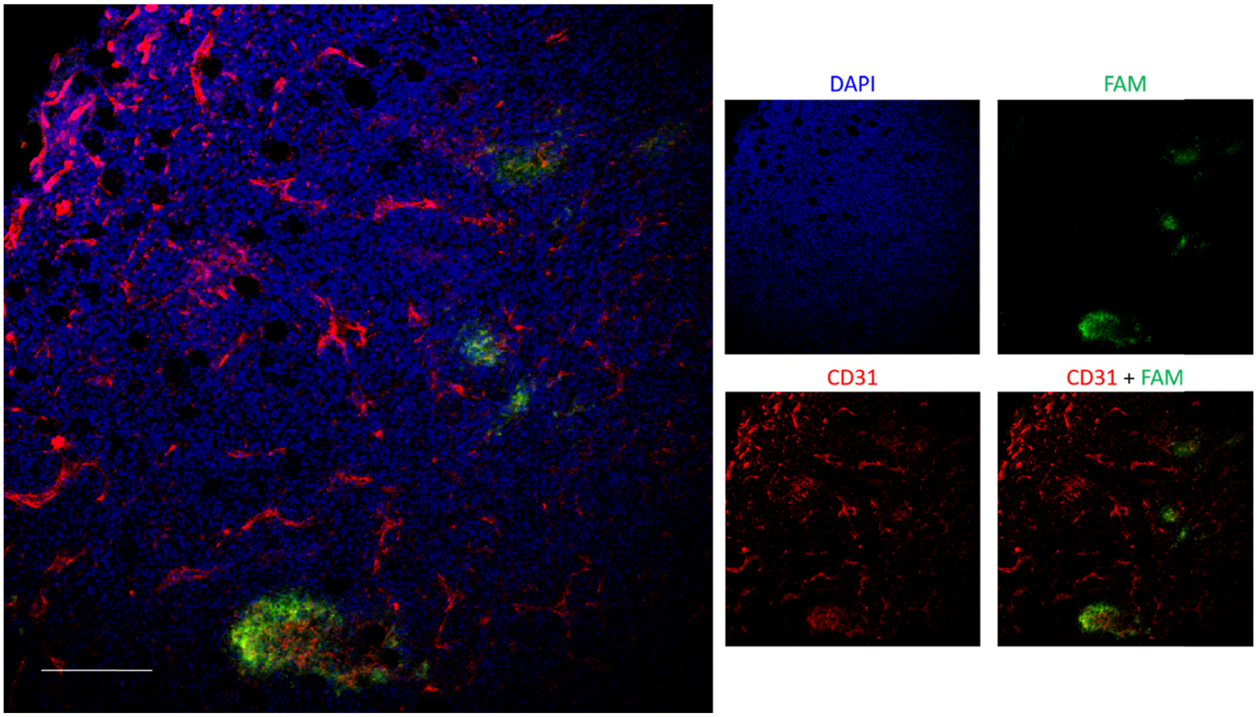


**Figure S10.** Tumor homing of PEGylated $CL_{PTX}$ vector containing 5 mol% PEG5K-lipid assessed by immunofluorescence microscopy. The formulation was administered intravenously in 4T1 tumor-bearing mouse; 24 h later the mouse was perfused with PBS, tumor excised, cryo-sectioned, and immunostained for CD31 (blood vessels; red) and stained with DAPI nuclear counterstain (blue); the green signal represents the FAM fluorescence of the PEG-$CL_{PTX}$. Tumor from mouse #6. Scale bar: 200 µm.

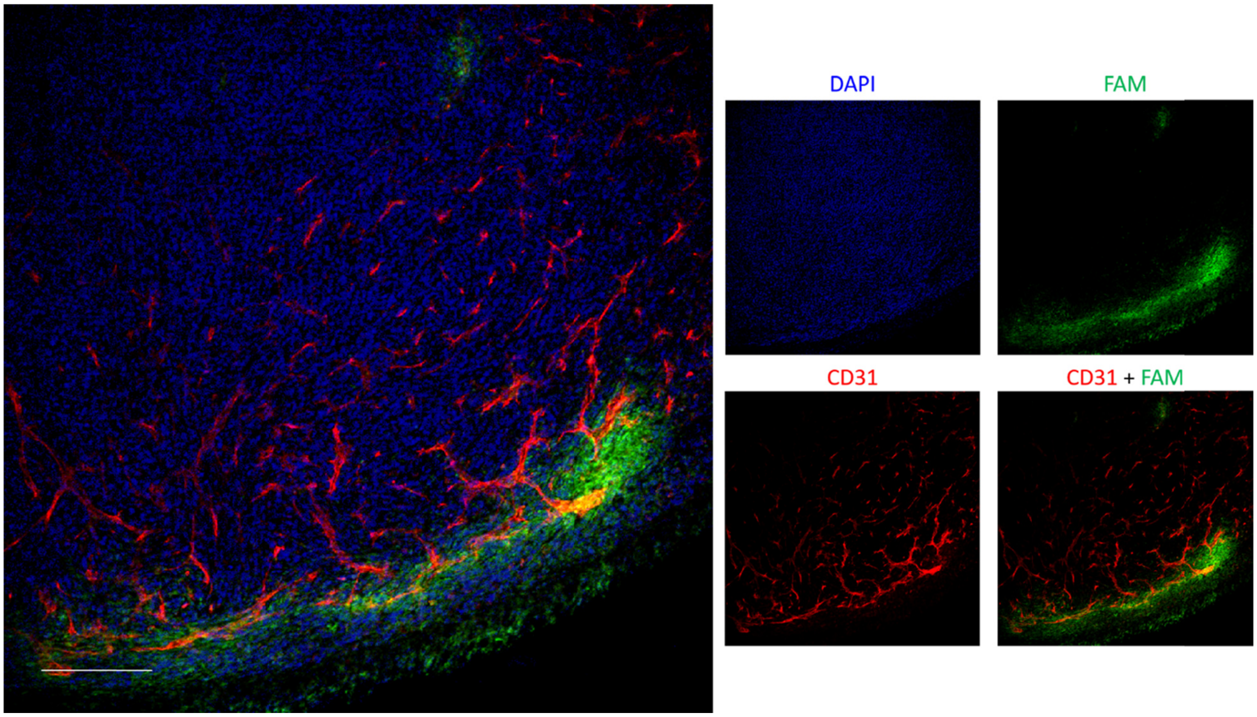


**Figure S11.** Tumor homing of PEGylated $CL_{PTX}$ vector containing 10 mol% PEG5K-lipid assessed by immunofluorescence microscopy. The formulation was administered intravenously in 4T1 tumor-bearing mouse; 24 h later the mouse was perfused with PBS, tumor excised, cryo-sectioned, and immunostained for CD31 (blood vessels; red) and stained with DAPI nuclear counterstain (blue); the green signal represents the FAM fluorescence of the PEG-$CL_{PTX}$. Tumor from mouse #7. Scale bar: 200 µm.

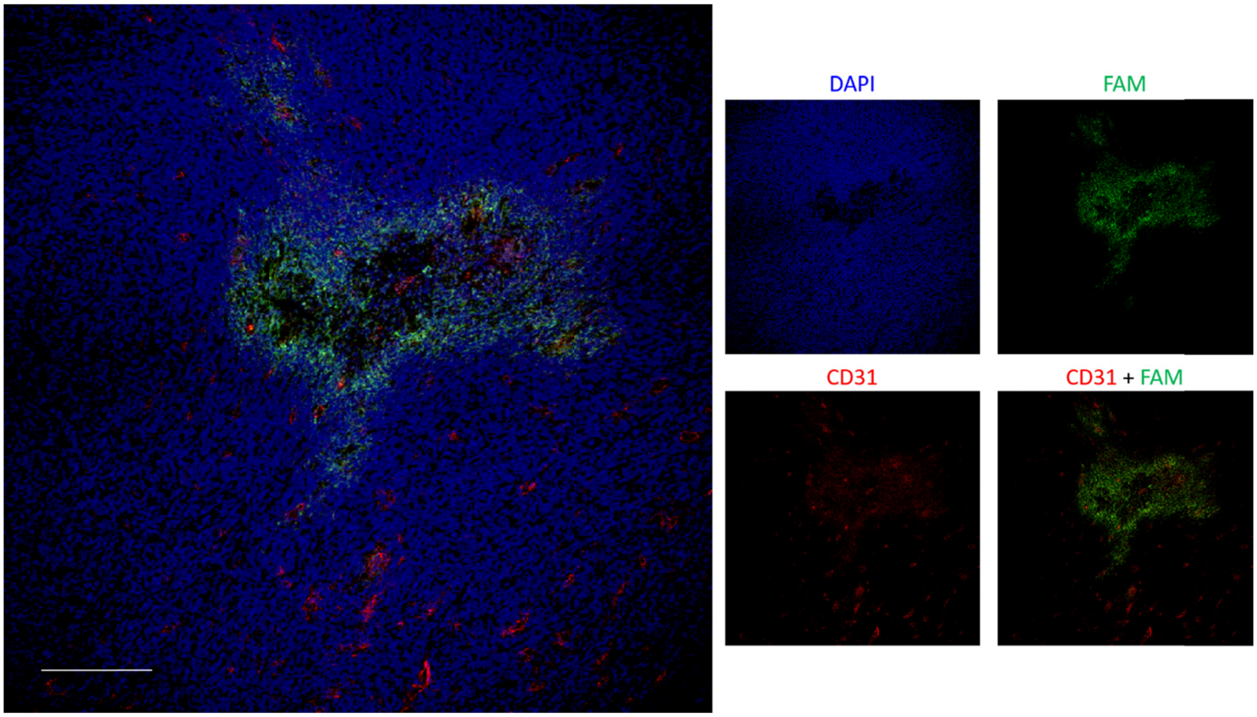


**Figure S12.** Tumor homing of PEGylated $CL_{PTX}$ vector containing 10 mol% PEG5K-lipid assessed by immunofluorescence microscopy. The formulation was administered intravenously in 4T1 tumor-bearing mouse; 24 h later the mouse was perfused with PBS, tumor excised, cryo-sectioned, and immunostained for CD31 (blood vessels; red) and stained with DAPI nuclear counterstain (blue); the green signal represents the FAM fluorescence of the PEG-$CL_{PTX}$. Tumor from mouse #8. Scale bar: 200 µm.

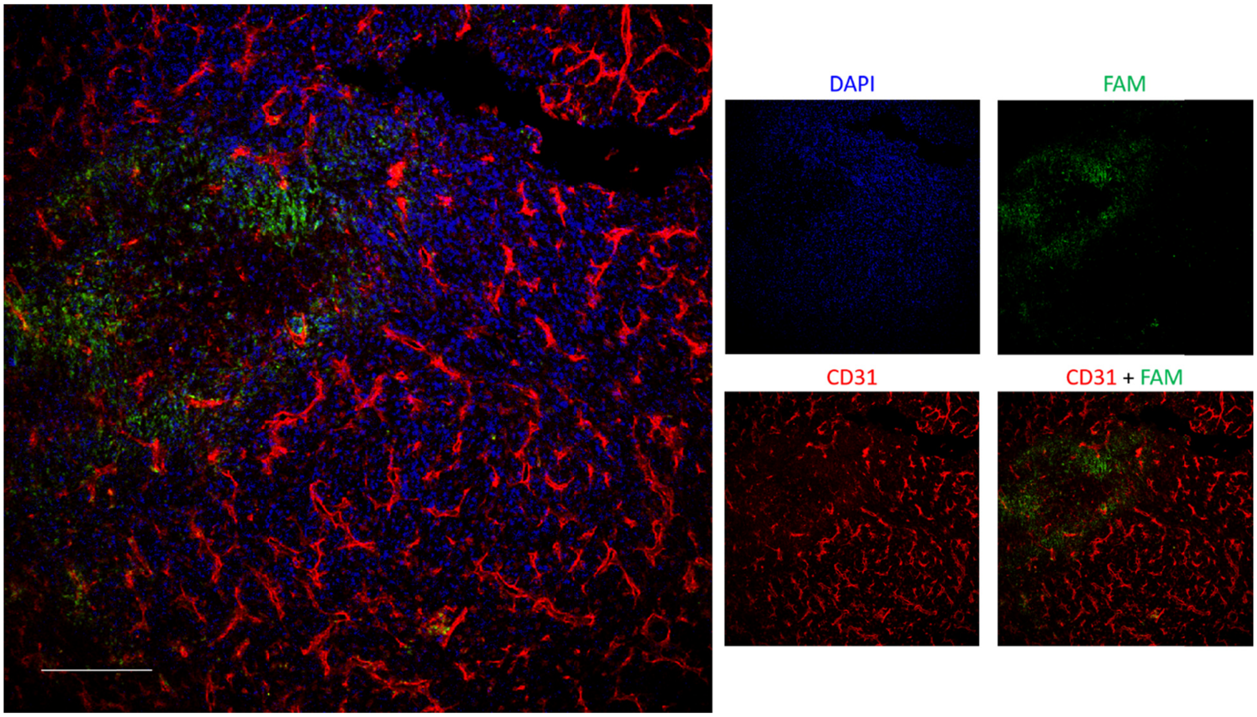


**Figure S13.** Tumor homing of PEGylated $CL_{PTX}$ vector containing 10 mol% PEG5K-lipid assessed by immunofluorescence microscopy. The formulation was administered intravenously in 4T1 tumor-bearing mouse; 24 h later the mouse was perfused with PBS, tumor excised, cryo-sectioned, and immunostained for CD31 (blood vessels; red) and stained with DAPI nuclear counterstain (blue); the green signal represents the FAM fluorescence of the PEG-$CL_{PTX}$. Tumor from mouse #9. Scale bar: 200 µm.

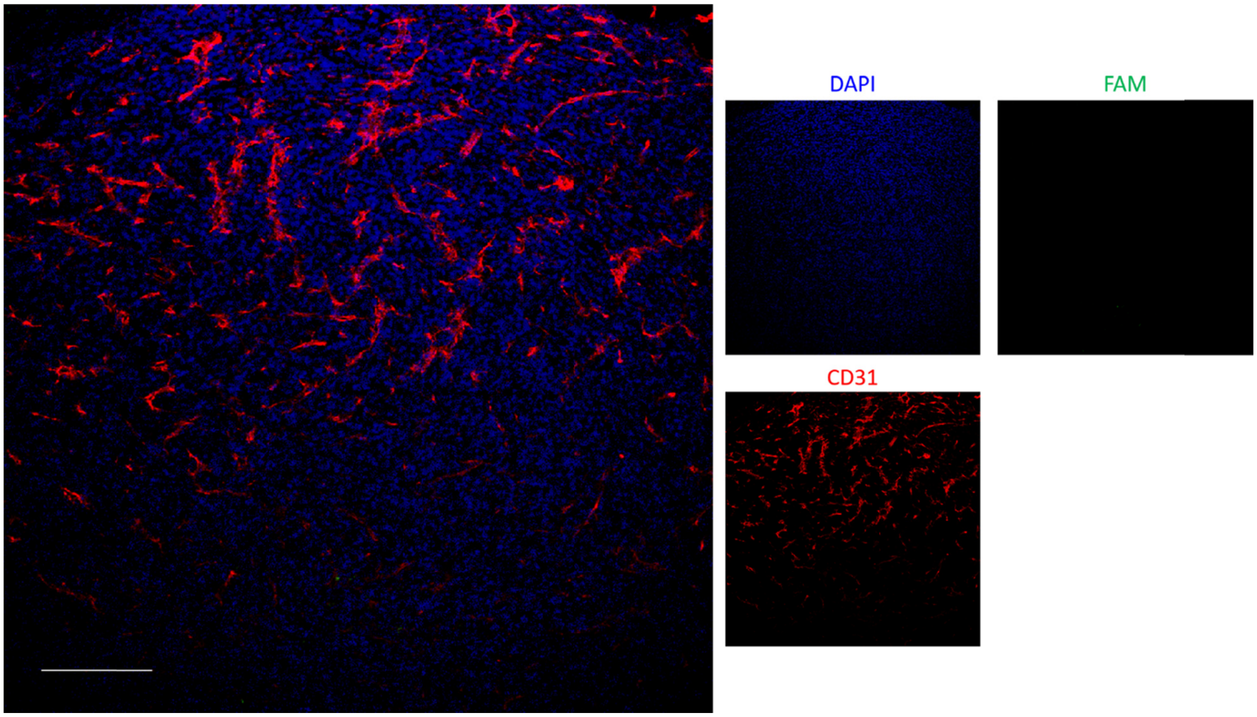


**Figure S14.** Immunofluorescence microscopy images of 4T1 tumor sections of a nontreated mouse. The mouse was perfused with PBS, tumor excised, cryo-sectioned, and immunostained for CD31 (blood vessels; red) and stained with DAPI nuclear counterstain (blue); the green signal represents the FAM fluorescence of the PEG-$CL_{PTX}$. Tumor from mouse #10. Scale bar: 200 µm.

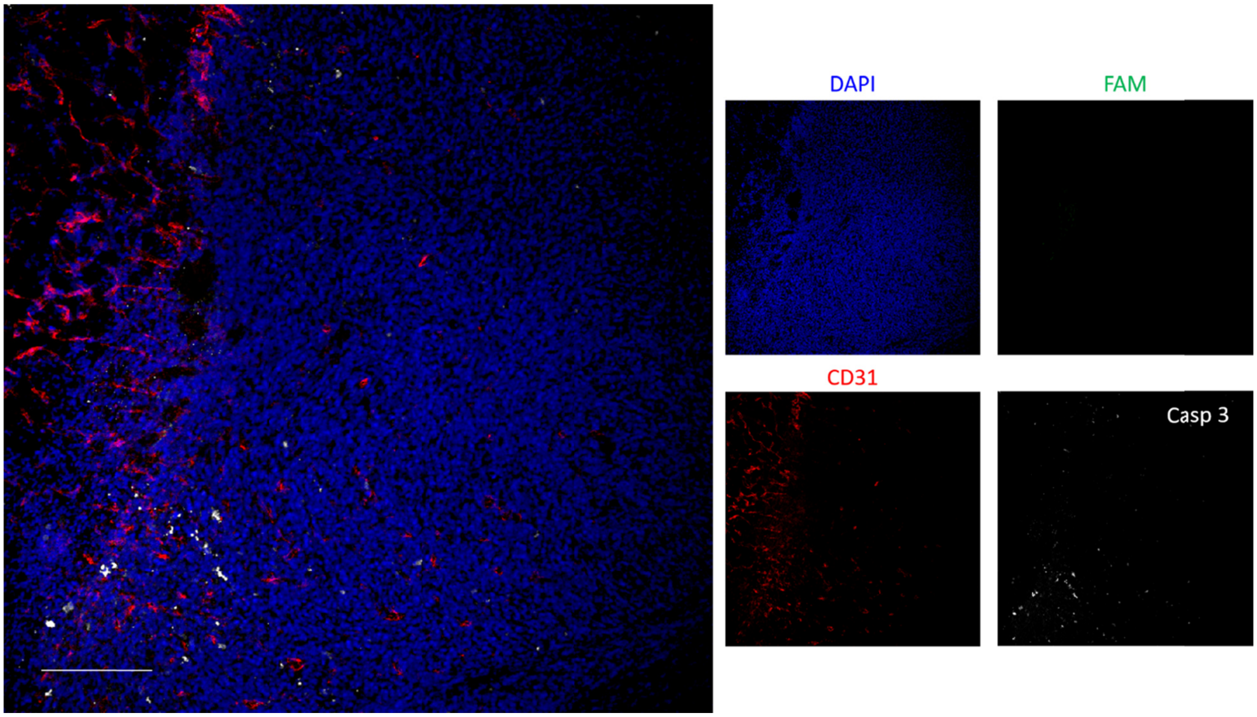


**Figure S15.** Tumor homing and cell death of PEGylated $CL_{PTX}$ vector containing 2 mol% PEG5K-lipid assessed by immunofluorescence microscopy. The formulation was administered intravenously in 4T1 tumor-bearing mouse; 24 h later the mouse was perfused with PBS, tumor excised, cryo-sectioned, and immunostained for CD31 (blood vessels; red) and cleaved caspase-3 (white), and stained with DAPI nuclear counterstain (blue); the green signal represents the FAM fluorescence of the PEG-$CL_{PTX}$. Tumor from mouse #1. Scale bar: 200 μm.

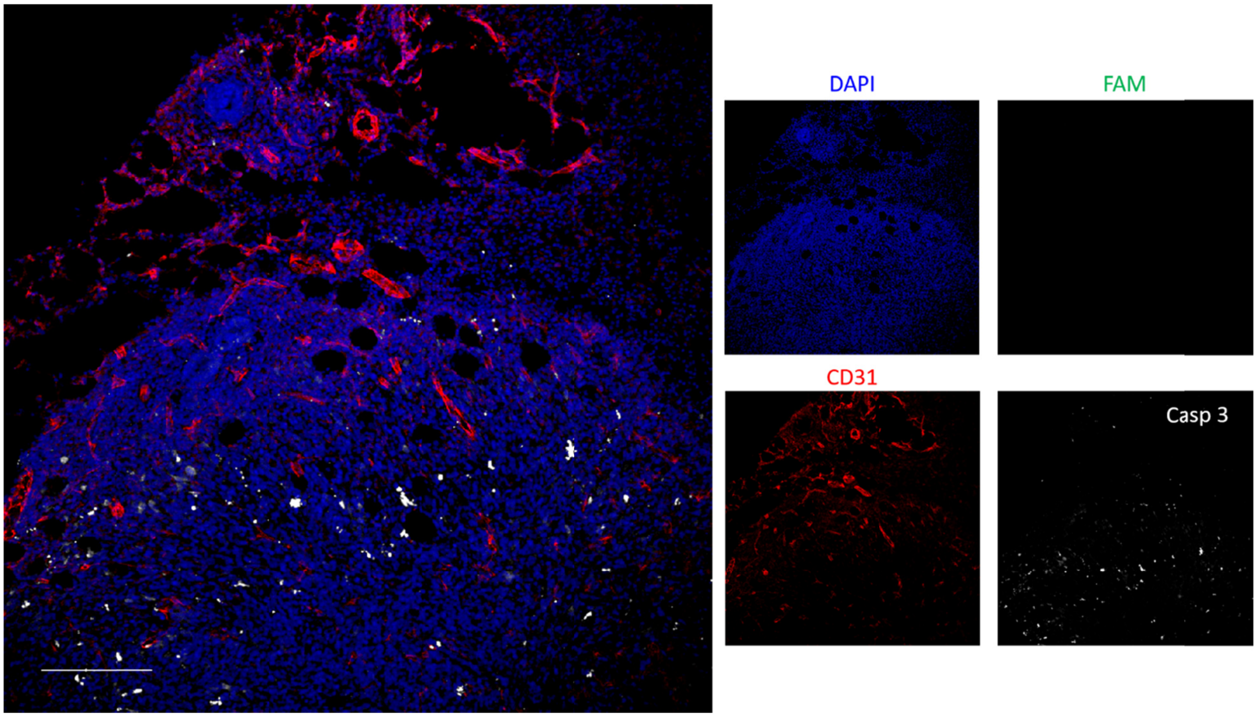


**Figure S16.** Tumor homing and cell death of PEGylated $CL_{PTX}$ vector containing 2 mol% PEG5K-lipid assessed by immunofluorescence microscopy. The formulation was administered intravenously in 4T1 tumor-bearing mouse; 24 h later the mouse was perfused with PBS, tumor excised, cryo-sectioned, and immunostained for CD31 (blood vessels; red) and cleaved caspase-3 (white), and stained with DAPI nuclear counterstain (blue); the green signal represents the FAM fluorescence of the PEG-$CL_{PTX}$. Tumor from mouse #2. Scale bar: 200 μm.

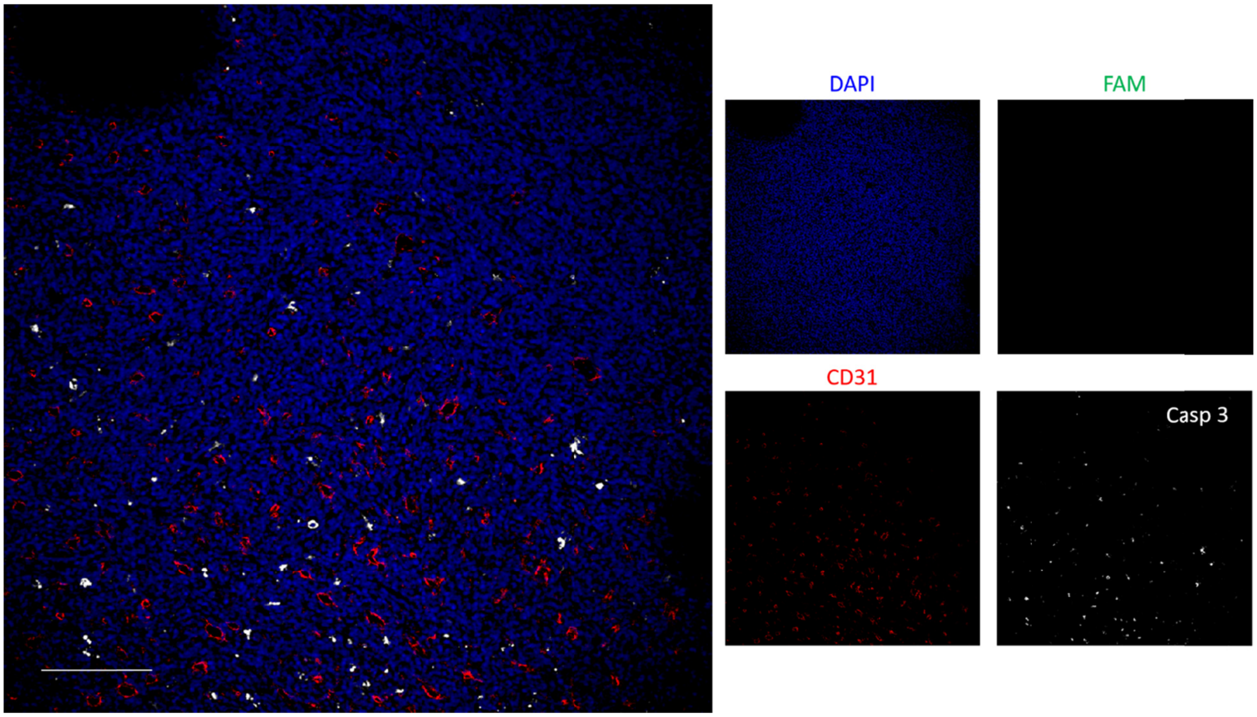


**Figure S17.** Tumor homing and cell death of PEGylated $CL_{PTX}$ vector containing 2 mol% PEG5K-lipid assessed by immunofluorescence microscopy. The formulation was administered intravenously in 4T1 tumor-bearing mouse; 24 h later the mouse was perfused with PBS, tumor excised, cryo-sectioned, and immunostained for CD31 (blood vessels; red) and cleaved caspase-3 (white), and stained with DAPI nuclear counterstain (blue); the green signal represents the FAM fluorescence of the PEG-$CL_{PTX}$. Tumor from mouse #3. Scale bar: 200 μm.

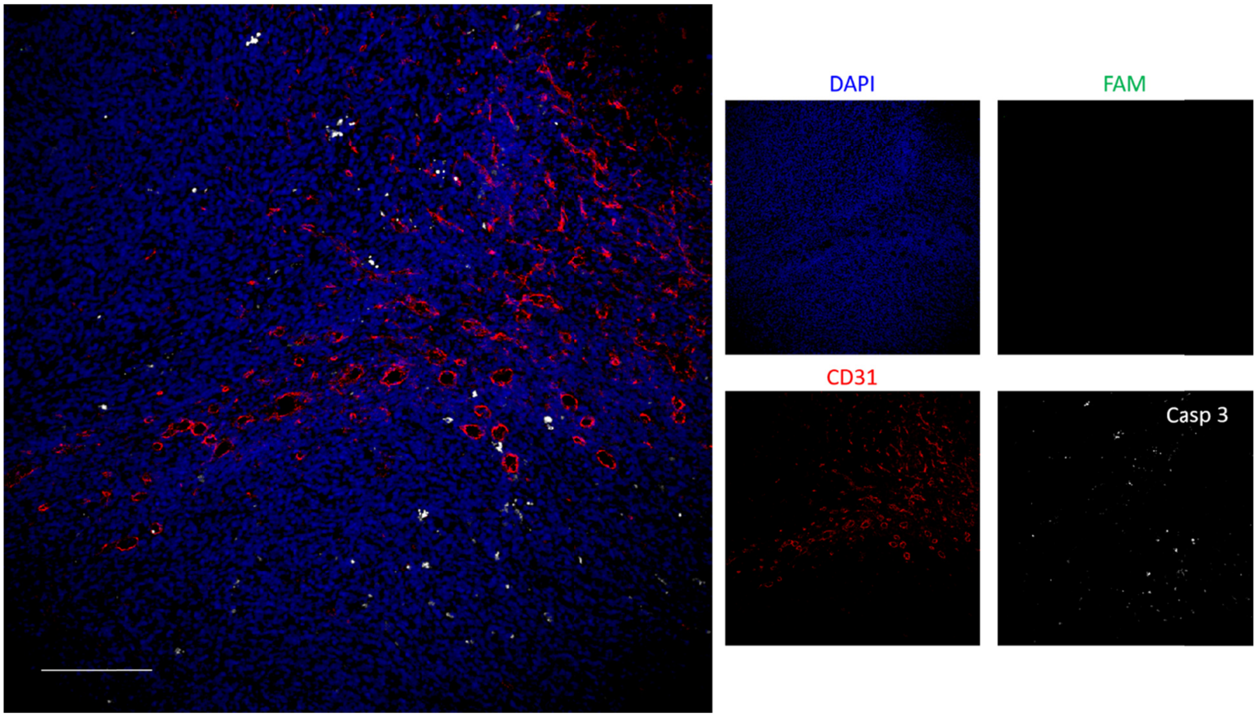


**Figure S18.** Tumor homing and cell death of PEGylated $CL_{PTX}$ vector containing 5 mol% PEG5K-lipid assessed by immunofluorescence microscopy. The formulation was administered intravenously in 4T1 tumor-bearing mouse; 24 h later the mouse was perfused with PBS, tumor excised, cryo-sectioned, and immunostained for CD31 (blood vessels; red) and cleaved caspase-3 (white), and stained with DAPI nuclear counterstain (blue); the green signal represents the FAM fluorescence of the PEG-$CL_{PTX}$. Tumor from mouse #4. Scale bar: 200 µm.

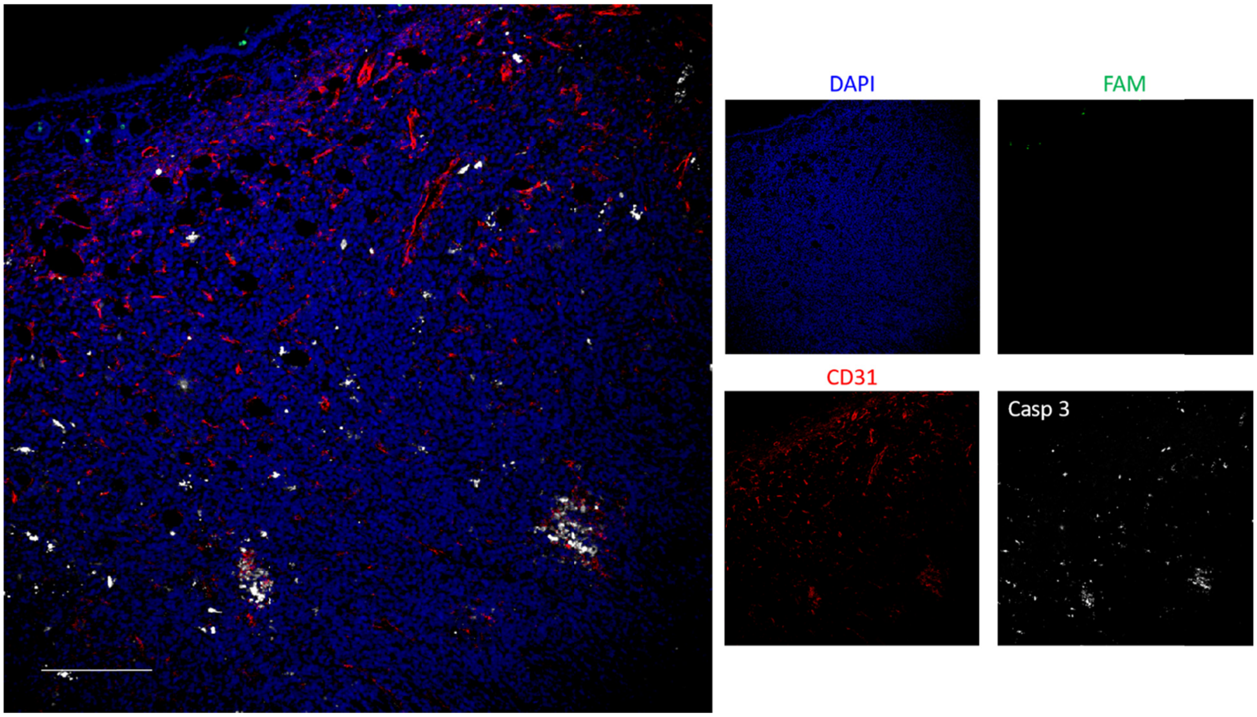


**Figure S19.** Tumor homing and cell death of PEGylated $CL_{PTX}$ vector containing 5 mol% PEG5K-lipid assessed by immunofluorescence microscopy. The formulation was administered intravenously in 4T1 tumor-bearing mouse; 24 h later the mouse was perfused with PBS, tumor excised, cryo-sectioned, and immunostained for CD31 (blood vessels; red) and cleaved caspase-3 (white), and stained with DAPI nuclear counterstain (blue); the green signal represents the FAM fluorescence of the PEG-$CL_{PTX}$. Tumor from mouse #5. Scale bar: 200 μm.

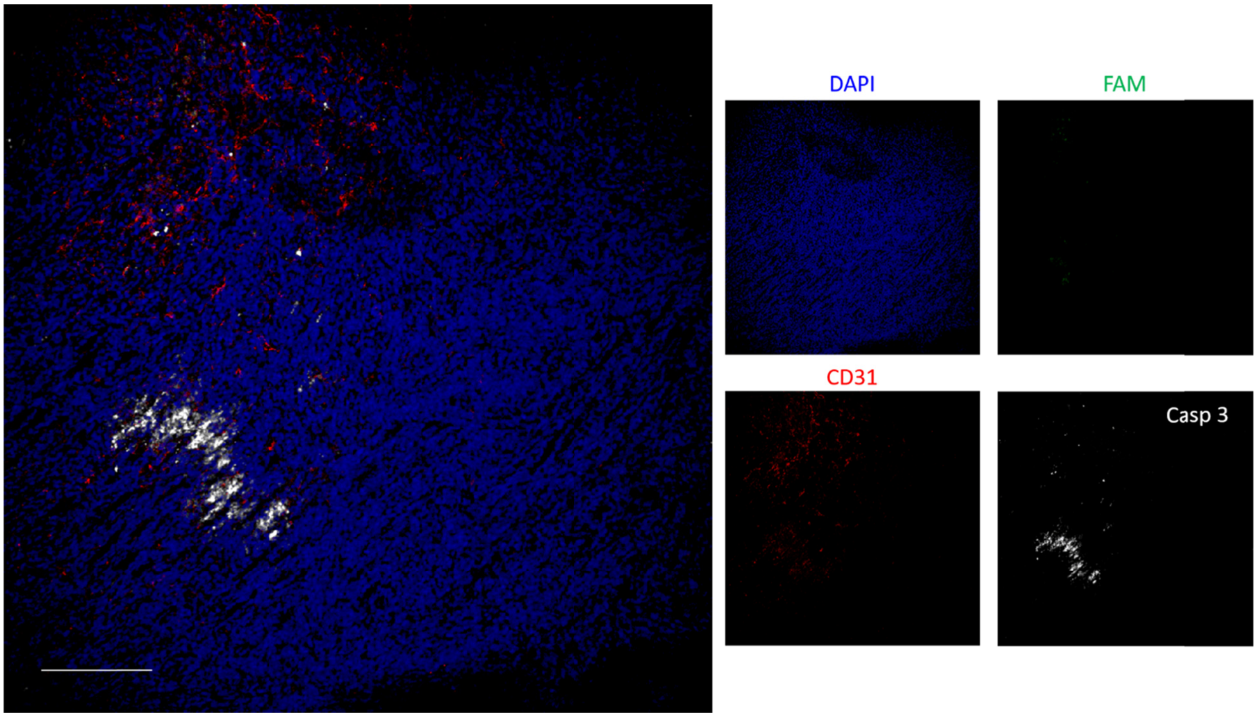


**Figure S20.** Tumor homing and cell death of PEGylated $CL_{PTX}$ vector containing 5 mol% PEG5K-lipid assessed by immunofluorescence microscopy. The formulation was administered intravenously in 4T1 tumor-bearing mouse; 24 h later the mouse was perfused with PBS, tumor excised, cryo-sectioned, and immunostained for CD31 (blood vessels; red) and cleaved caspase-3 (white), and stained with DAPI nuclear counterstain (blue); the green signal represents the FAM fluorescence of the PEG-$CL_{PTX}$. Tumor from mouse #6. Scale bar: 200 µm.

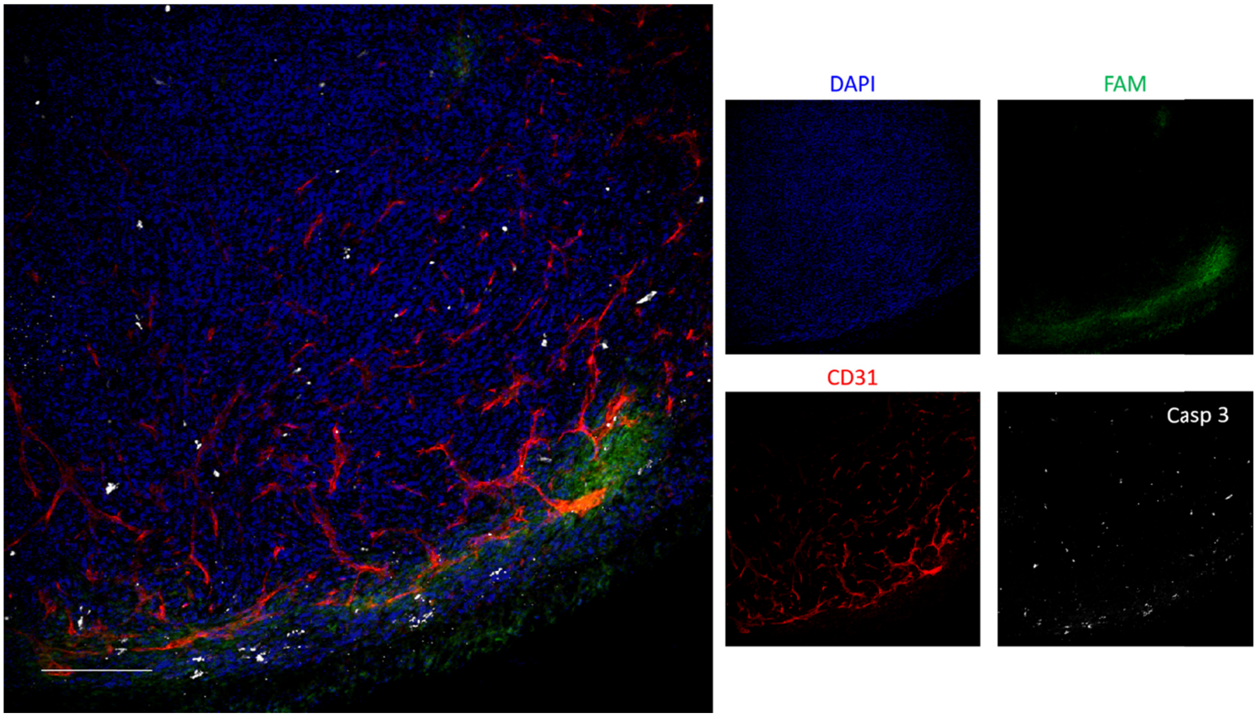


**Figure S21.** Tumor homing and cell death of PEGylated $CL_{PTX}$ vector containing 10 mol% PEG5K-lipid assessed by immunofluorescence microscopy. The formulation was administered intravenously in 4T1 tumor-bearing mouse; 24 h later the mouse was perfused with PBS, tumor excised, cryo-sectioned, and immunostained for CD31 (blood vessels; red) and cleaved caspase-3 (white), and stained with DAPI nuclear counterstain (blue); the green signal represents the FAM fluorescence of the PEG-$CL_{PTX}$. Tumor from mouse #7. Scale bar: 200 μm.

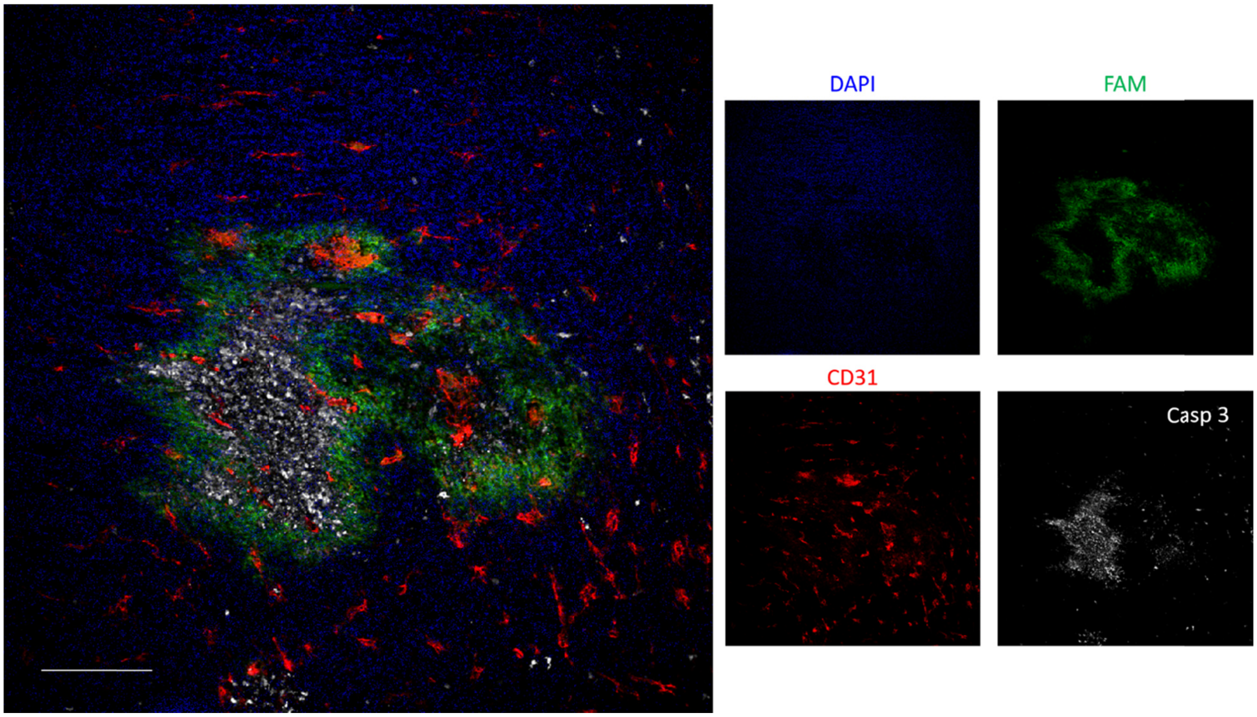


**Figure S22.** Tumor homing and cell death of PEGylated $CL_{PTX}$ vector containing 10 mol% PEG5K-lipid assessed by immunofluorescence microscopy. The formulation was administered intravenously in 4T1 tumor-bearing mouse; 24 h later the mouse was perfused with PBS, tumor excised, cryo-sectioned, and immunostained for CD31 (blood vessels; red) and cleaved caspase-3 (white), and stained with DAPI nuclear counterstain (blue); the green signal represents the FAM fluorescence of the PEG-$CL_{PTX}$. Tumor from mouse #8. Scale bar: 200 μm.

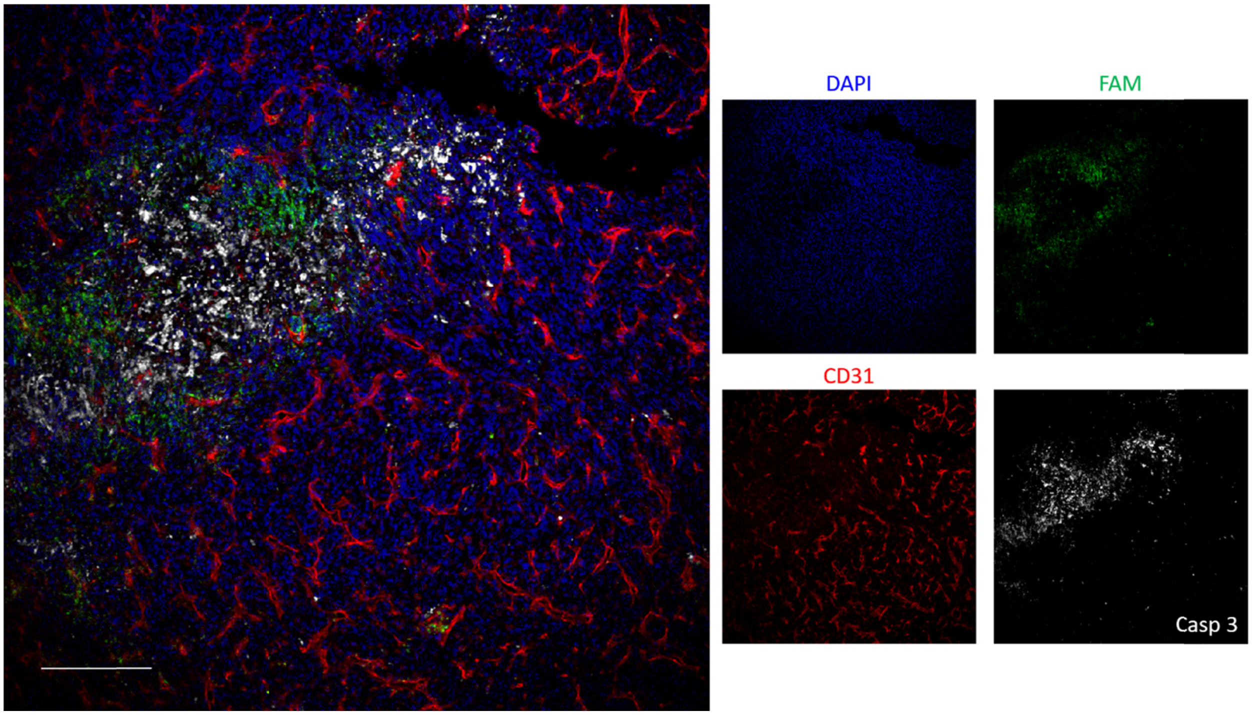


**Figure S23.** Tumor homing and cell death of PEGylated $CL_{PTX}$ vector containing 10 mol% PEG5K-lipid assessed by immunofluorescence microscopy. The formulation was administered intravenously in 4T1 tumor-bearing mouse; 24 h later the mouse was perfused with PBS, tumor excised, cryo-sectioned, and immunostained for CD31 (blood vessels; red) and cleaved caspase-3 (white), and stained with DAPI nuclear counterstain (blue); the green signal represents the FAM fluorescence of the PEG-$CL_{PTX}$. Tumor from mouse #9. Scale bar: 200 µm.

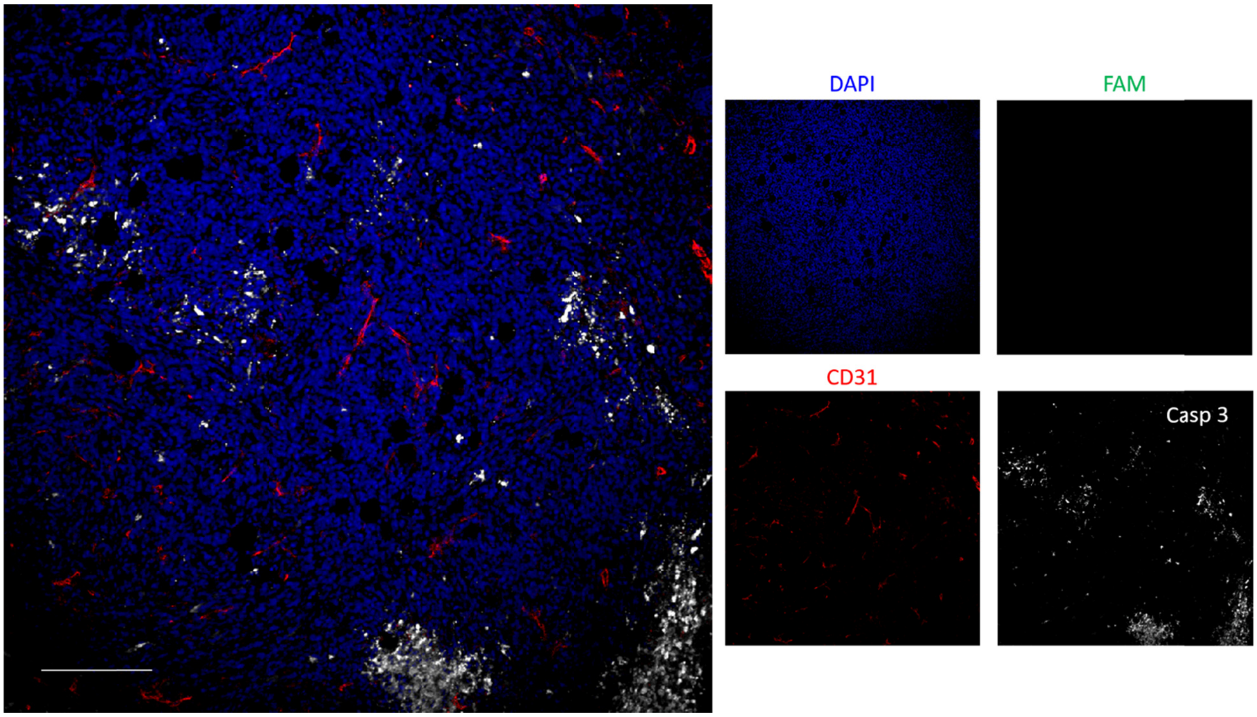


**Figure S24.** Tumor homing and cell death of PEGylated CL without PTX vector containing 10 mol% PEG5K-lipid assessed by immunofluorescence microscopy. The formulation was administered intravenously in 4T1 tumor-bearing mouse; 24 h later the mouse was perfused with PBS, tumor excised, cryo-sectioned, and immunostained for CD31 (blood vessels; red) and cleaved caspase-3 (white), and stained with DAPI nuclear counterstain (blue); the green signal represents the FAM fluorescence of the PEG-CL. Tumor from mouse #11. Scale bar: 200 μm.

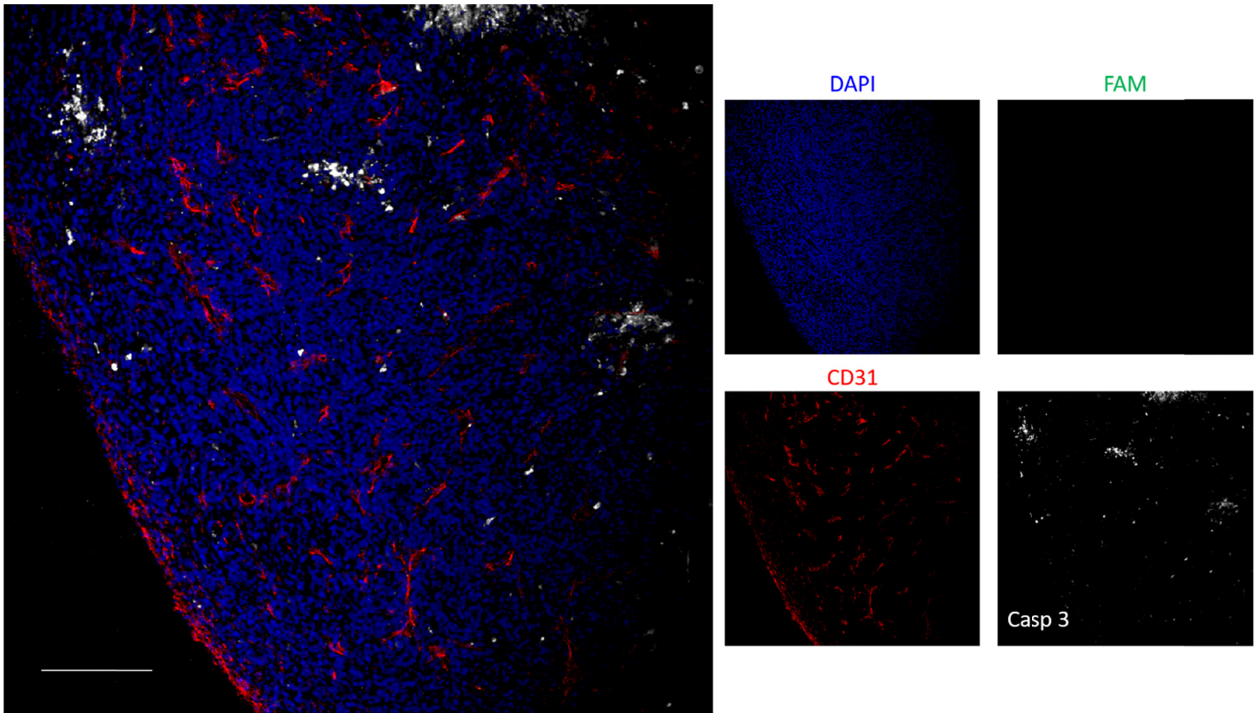


**Figure S25.** Tumor homing and cell death of PEGylated CL without PTX vector containing 10 mol% PEG5K-lipid assessed by immunofluorescence microscopy. The formulation was administered intravenously in 4T1 tumor-bearing mouse; 24 h later the mouse was perfused with PBS, tumor excised, cryo-sectioned, and immunostained for CD31 (blood vessels; red) and cleaved caspase-3 (white), and stained with DAPI nuclear counterstain (blue); the green signal represents the FAM fluorescence of the PEG-CL. Tumor from mouse #12. Scale bar: 200 µm.

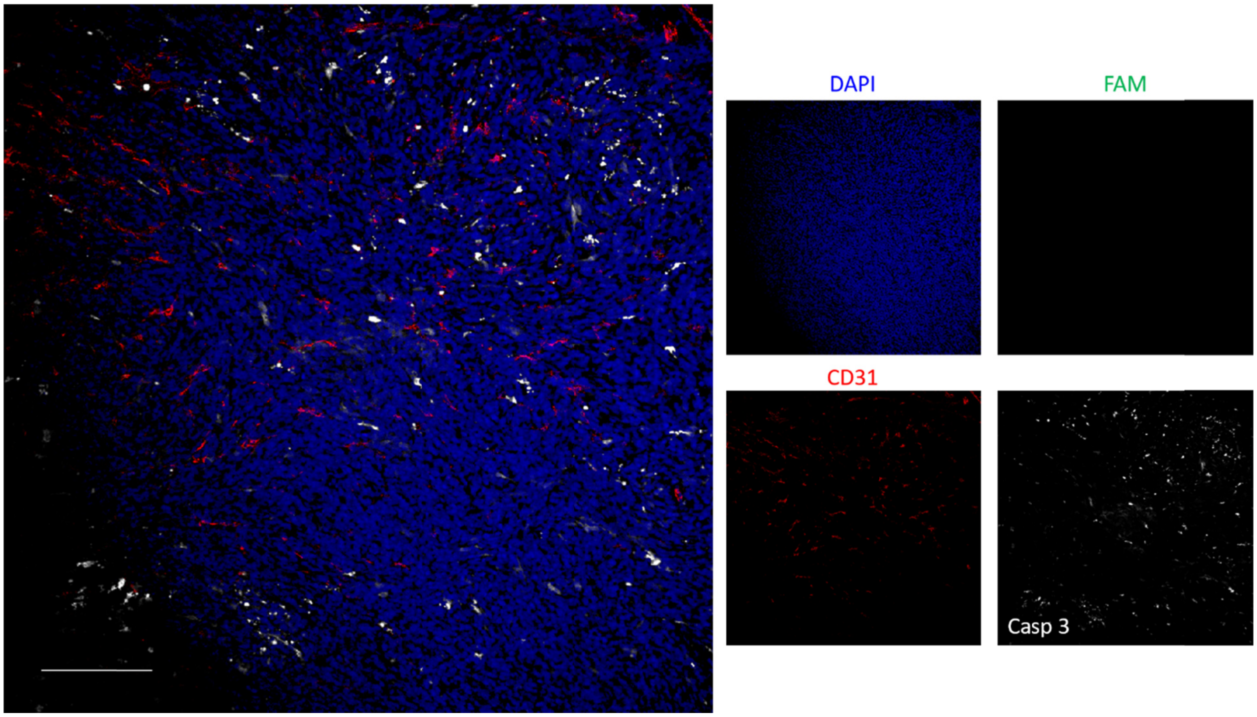


**Figure S26.** Tumor homing and cell death of PEGylated CL without PTX vector containing 10 mol% PEG5K-lipid assessed by immunofluorescence microscopy. The formulation was administered intravenously in 4T1 tumor-bearing mouse; 24 h later the mouse was perfused with PBS, tumor excised, cryo-sectioned, and immunostained for CD31 (blood vessels; red) and cleaved caspase-3 (white), and stained with DAPI nuclear counterstain (blue); the green signal represents the FAM fluorescence of the PEG-CL. Tumor from mouse #13. Scale bar: 200 µm.

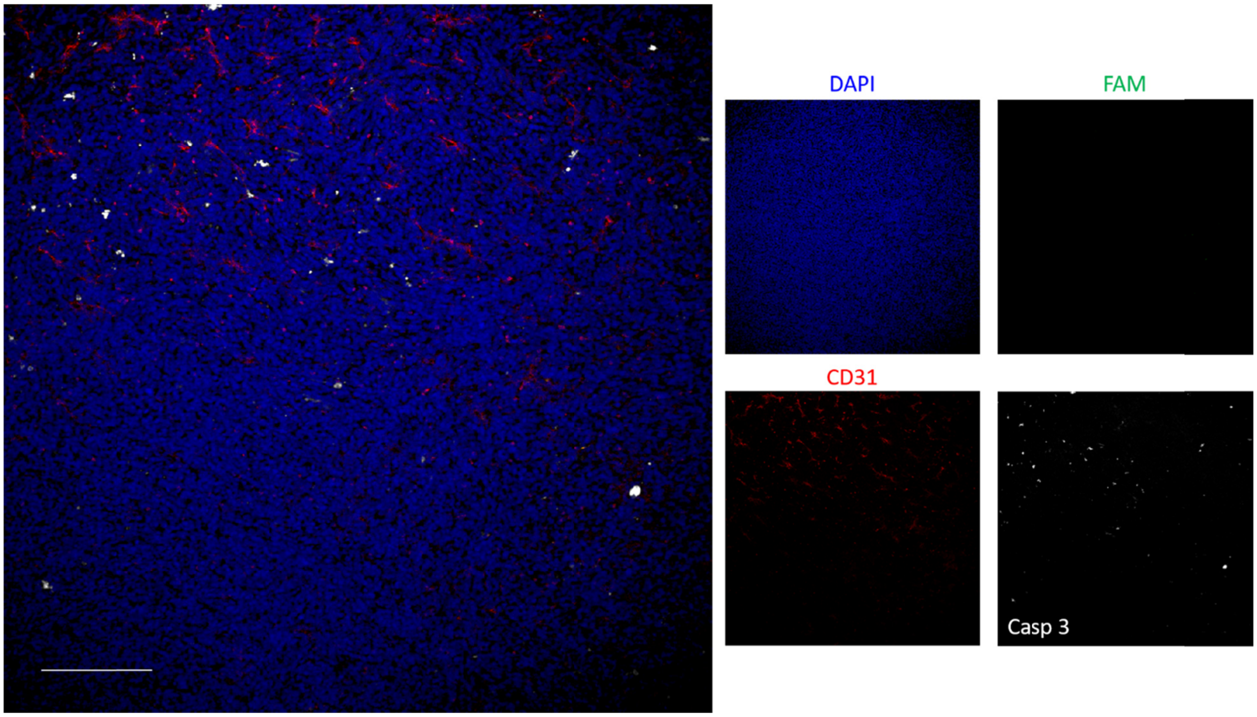


**Figure S27.** Immunofluorescence microscopy images of 4T1 tumor sections of a nontreated mouse. The mouse was perfused with PBS, tumor excised, cryo-sectioned, and immunostained for CD31 (blood vessels; red), and cleaved caspase-3 (white), and stained with DAPI nuclear counterstain (blue); the green signal represents the FAM fluorescence of the PEG-$CL_{PTX}$. Tumor from mouse #10. Scale bar: 200 µm.
